# The Combination of Morphology and Surface Chemistry Defines the Biological Identity of Nanocarriers in Human Blood

**DOI:** 10.1101/2020.09.02.280404

**Authors:** Nicholas B. Karabin, Michael P. Vincent, Sean D. Allen, Sharan Bobbala, Molly A. Frey, Sijia Yi, Yufan Yang, Evan A. Scott

## Abstract

Following intravenous administration, an adsorbed corona of blood proteins immediately forms on the surfaces of nanocarriers to confer a distinct biological identity that dictates interactions with the immune system. While the nanocarrier surface chemistry has long been the focus of protein corona formation, the influence of the nanocarrier structure has remained unclear despite well-documented influences on biodistribution, clearance and inflammation. Here, we present design rules for the combined engineering of both nanocarrier structure and surface chemistry derived from a comprehensive proteomic analysis of protein corona formation in human blood. A library of nine soft PEGylated nanocarriers that differ in their combination of morphology (spheres, vesicles, and cylinders) and surface chemistry (methoxy, hydroxyl, and phosphate) were synthesized to represent properties of commonly employed drug delivery vehicles. Using label-free proteomics and high-throughput techniques, we examined the relationship between physicochemical properties and the resulting nanocarrier biological identity, including dynamic changes in protein corona composition, differential immunostimulation and uptake by relevant immune cell populations. In human blood, non-polar spherical micelles developed a similar biological identity to polar vesicles, whereas the identities of polar spheres and cylinders resembled that of non-polar vesicles. The formed protein coronas were compositionally dynamic and morphology-dependent, and these time-dependent fingerprints altered nanocarrier complement activation as well as their uptake by human monocytes, macrophages, and dendritic cells. This comprehensive analysis provides mechanistic insights into rational design choices that impact nanocarrier fate in human blood.

**One Sentence Summary:** We demonstrate that not only the surface chemistry, but the combined chemical and structural properties of soft drug delivery vehicles impact the composition of blood proteins that adsorb to their surfaces, and these differences specify their interactions with and modulation of human immune cells.

## Introduction

Nanoscale carriers, i.e. nanocarriers, are customizable delivery vehicles for therapeutic and diagnostic agents, and their fate following *in vivo* administration is highly dependent on their interactions with cells of the immune system (*1*), and are of significant interest for applications within the immunomodulation space (*2-4*). In these applications, nanocarriers are utilized with the intent of altering the host’s immune system by eliciting either a pro-or anti-inflammatory response through targeting various immune cell subsets. While the cellular target may vary based on the specific application or disease state, the cells of the mononuclear phagocyte system (MPS), namely dendritic cells, macrophages, and monocytes, have garnered significant attention due to their importance in maintaining homeostasis and contributions to inflammation and autoimmunity (*5*). A paradoxical scenario is presented by the diverse cellular functions of the MPS, where these cells represent both desirable targets for the administered therapy, as well as sources of nonspecific uptake capable of quickly clearing the nanocarrier from circulation before it can complete its desired function (*4, 6*). As such, understanding how nanocarriers can be rationally designed to avoid or target specific components of the MPS has been an area of active investigation. These efforts have included both active (*7*) and passive (*8, 9*) targeting strategies that typically involve the respective surface display of targeting ligands or the modulation of either surface chemistry or morphology of the nanocarrier (*10-12*).

After introducing nanocarriers into the biological milieu, the once meticulously designed and characterized synthetic identity of the nanocarrier surface is immediately replaced by a new biological identity defined by the adsorbed protein corona (*13*). This dynamic shell of adsorbed biomolecules at the nanocarrier-fluid interface is the actual surface with which cells directly interact and, as such, plays a central role is defining the cellular fate of the nanocarrier (*14*). A variety of physicochemical characteristics, including nanocarrier morphology (*15*), surface chemistry (*16, 17*), surface charge (*17, 18*), and porosity (*19*), influence both the amount and identity of the absorbed protein. These sometimes subtle compositional changes within the protein corona have been attributed to both the promotion (*20*) and suppression (*21*) of nanocarrier internalization by MPS cells. These observations suggest that rationally designing the nanocarrier chassis (*i.e.* combined chemical and structural framework) to govern the composition of the protein corona, rather than to prevent its formation, is a potential strategy for tailoring cellular uptake.

Poly(ethylene glycol)-*block*-poly(propylene sulfide) (PEG-*b*-PPS) is an established block copolymer (BCP) that has been employed in the fabrication of a variety of nanocarrier chassis designed for immunomodulatory applications (*22-29*). The chassis morphology is dictated by the BCP composition and hydrophilic mass fraction (f_PEG_), allowing the formation of PEG-*b*-PPS spherical micelles (MCs) (*27*), vesicular polymersomes (PSs) (*24*), and cylindrical filomicelles (FMs) (*30*). For the PEG-*b*-PPS BCP system, morphological complexity is inversely related to f_PEG_, where f_PEG_ < 0.12 results in BCNs, 0.22 < f_PEG_ < 0.35 results in PSs, 0.37 < f_PEG_ < 0.39 results in FMs, and f_PEG_ > 0.45 forms MCs (*28, 31*). This characteristic of self-assembled systems allows for the preparation of diverse morphologies with equivalent chemical identities, and thereby permits the evaluation of morphology-driven variation in nanocarrier chassis performance. Previous studies have highlighted that PEG-*b*-PPS nanocarrier morphology is instrumental in governing material-host interactions, as evidenced by the altered organ-and cellular-level biodistributions achieved by these nanocarriers (*23, 32-34*). However, attempts to further tailor cellular interactions of PEG-*b*-PPS nanocarriers by combining the influence of morphology with surface chemistry, have been limited (*35*).

Here, we investigate how combinations of surface chemistry and morphology can be used to influence PEG-*b*-PPS nanocarrier uptake within the primary cell subsets of the MPS. Previous studies involving PEG-*b*-PPS and numerous other PEGylated nanocarriers and drugs have largely utilized methoxy (MeO) or hydroxy (OH)-functionalization. In this work, we additionally describe the synthesis of two biomimetic phosphate (Phos)-terminated PEG-*b*-PPS BCPs that self-assemble into micelles and filomicelles. We previously described the synthesis of a phosphate-terminated PEG-*b*-PPS BCP that self assembles into PS (*17*). These nine BCPs result in a total of nine nanocarrier formulations that vary in their combination of morphology (sphere, vesicle and cylinder) and surface chemistry (MeO, OH, and Phos). We employ this soft nanocarrier library to systematically evaluate material rational design choices on the biological performance nanocarrier chassis in human blood. We envision the information gained from these studies will prove invaluable to the future design and development of high performing soft nanocarriers that have optimal biological identities for specific applications of interest (drug delivery, vaccination, etc.).

We apply a holistic *in vitro* approach that assesses protein adsorption, complement activation, and cytokine responses after introducing nanocarriers to human blood samples. To understand whether the composition of the protein corona is a function of combined physicochemical properties, we used modern label-free proteomics techniques to characterize the identity and relative abundance of human plasma proteins that define each protein corona at multiple timepoints (**Fig. 1A**). After conducting a thorough systems-level analysis of differentially adsorbing proteins, we examined the degree to which each biological identity evokes opsonization and cytokine responses in the blood (**Fig. 1B**). We conclude our study by characterizing how various combinations of morphology and surface chemistry influence nanocarrier uptake by human monocytes, macrophages, and dendritic cells, which are the primary components of the MPS(**Fig. 1C**). We assess whether the differential uptake of the nine formulations can be understood in the context of their time-dependent adsorbed protein fingerprint. To the best of our knowledge, this work represents the most comprehensive demonstration and analysis of the link between soft nanocarrier physicochemical properties and human protein corona formation and the resulting consequences on nanocarrier interactions with human cells.

**Fig. 1.**
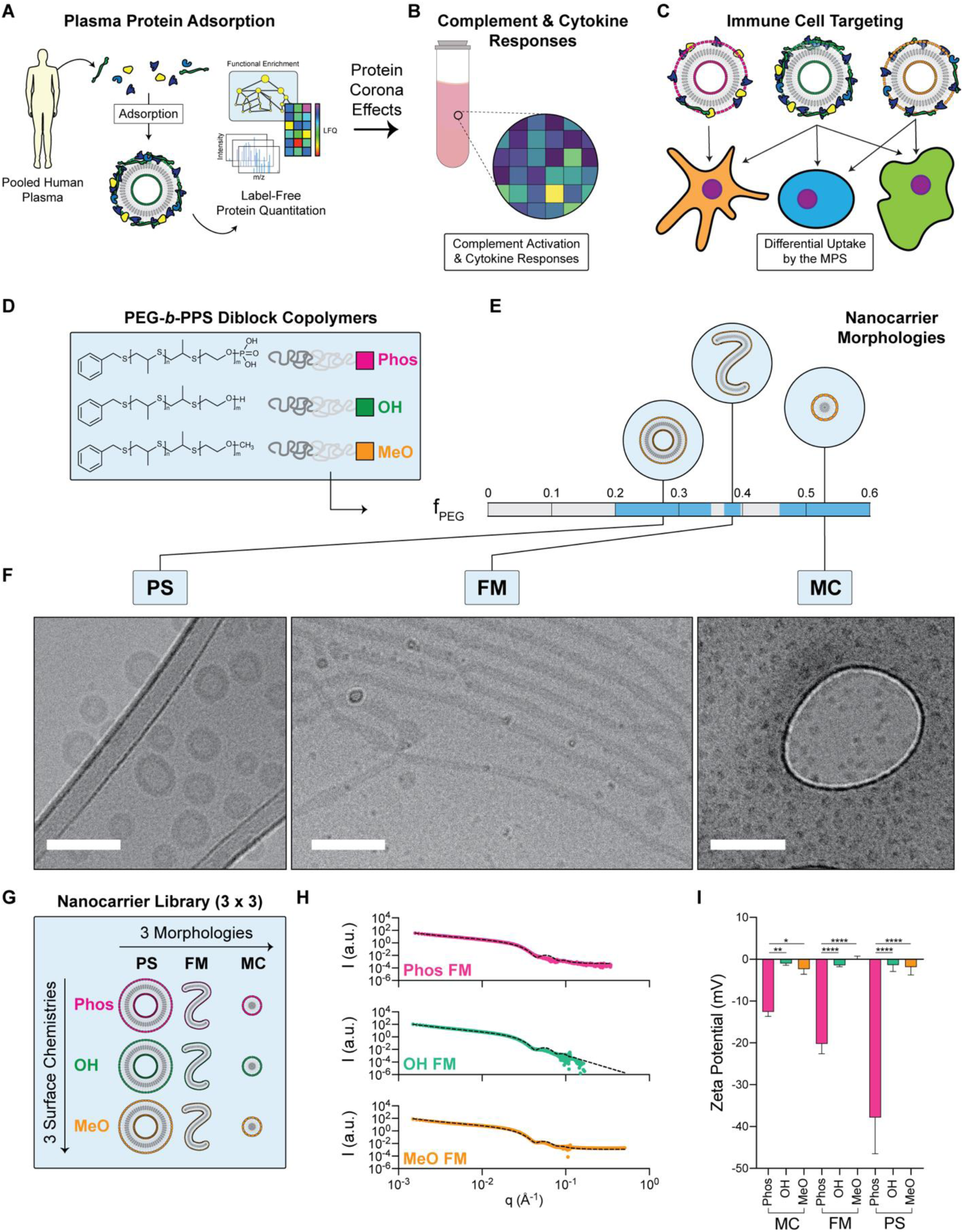
PEG-*b*-PPS nanocarrier library development, physicochemical characterization and overview of performance assessments. (**A-C**) Illustrative overview of the blood protein adsorption analysis and the consequence of formed protein coronas on nanocarrier performance. All studies use human derived biological samples. (**A**) The adsorption of human plasma proteins to soft nanocarrier surfaces were quantified used modern biochemical and label-free proteomics techniques. Each nanocarrier protein corona was characterized at 2 h and 24 h to assess the Vroman effect. The effect of the protein corona on (**B**) nanocarrier-induced complement activation and cytokine responses in human blood, and (**C**) nanocarrier uptake by the cells of the human mononuclear phagocyte system (MPS) was examined. (**D**) Schematic of the different functionalized BCPs terminated with phosphate-, hydroxyl-, or methoxy-chemical groups. (**E**) The BCP hydrophilic mass fraction (fPEG) was varied to promote the favorable self-assembly of three morphologies: polymersomes (PS), filomicelles (FM), and micelles (MC). (**F**) Cryogenic TEM micrographs of PS, FM, and MC nanocarrier morphologies (Scale bar = 100 nm). The OH-functionalized examples are displayed as an example. The complete morphological characterization is presented in **Figure S4**. (**G**) Illustration of the complete library of nanocarriers that differ in their combination of morphology and surface chemistry. (**H**) SAXS profiles of PEG-*b*-PPS FMs overlaid with corresponding fit for a flexible cylinder model (χ2 = 0.004, 0.0439, 0.777 for MeO-, OH-, and Phos-FMs, respectively). (**I**) Zeta potential of each nanocarrier formulation (n = 3). Significance was determined with Tukey’s multiple comparison test. \**p*<0.05, \*\**p*<0.01, \*\*\*\**p*<0.0001. Error bars represent s.d.

## Results

### Physicochemical Characterization of the PEG-b-PPS Nanocarrier Library

Phos-, OH-, and MeO-functionalized BCPs were each synthesized with varying f_PEG_ to prepare PSs, FMs, and MCs exhibiting each of the three aforementioned surface chemistries (**Fig. 1, D and E**). The reaction scheme used to prepare the BCPs utilized in this study is depicted by **Fig. S1**. All BCPs were characterized via NMR (**Fig. S2**) and gel permeation chromatography (**Fig. S3**). Phos-functionalized BCPs were further characterized using ^31^P NMR (**Fig. S2B**). The rightward shift of the phosphorous peak toward 0 ppm following hydrolysis indicated the successful formation of a monoester product (*36*). PSs, FMs, and MCs exhibiting each of the three surface chemistries were successfully prepared and characterized by cryogenic transmission electron microscopy (cryoTEM), dynamic light scattering (DLS), small angle x-ray scattering (SAXS), and electrophoretic light scattering (ELS). CryoTEM micrographs (**Fig. 1F**; **Fig. S4**) confirmed that each BCP, irrespective of the terminal group extending off the PEG block, self-assembled into its anticipated morphology as predicted by its f_PEG_.

DLS and SAXS were completed to assess the size characteristics of each nanocarrier in the library (**Fig. 1G**). DLS analysis of the spherical nanocarrier formulations indicated that PSs exhibited diameters ranging from 55 nm to 68 nm with polydispersity indices (PDIs) all below 0.150 (**Table S1**; **Fig. S4**). MCs ranged in diameter from 16 nm to 19 nm with PDIs below 0.120 (**Table S1**; **Fig. S4**). The morphology of each nanocarrier was further confirmed through SAXS (**Fig. S5**). Due to the anisotropic nature of FMs, size characteristics were solely derived from SAXS analysis. The SAXS scattering profile from each of the FM samples was fit utilizing a flexible cylinder model (**Fig. 1H**). Model fitting indicated that FMs exhibited core radii of 17 nm – 18 nm and contour lengths extending into the micron range (**Table S1**).

Lastly, the zeta potential of each nanocarrier formulation was assessed via ELS (**Fig. 1I**). PSs, FMs, and MCs exhibiting either hydroxyl or methoxy surface functionalities displayed zeta potentials ranging from-2 to 0 mV while those composed of Phos-functionalized BCPs displayed zeta potentials ranging from-38 to-13 mV. For nanocarriers composed of the Phos-functionalized BCPc, zeta potential measurements increased with the f_PEG_ of the BCPs (i.e. the smaller the f_PEG_ the more negative the zeta potential). There are several factors potentially contributing to these observed differences in the Phos-functionalized nanocarriers. First, differences in the molecular packing exhibited by the three morphologies results in variations in the surface area per BCP (*37*), which would manifest as a difference in zeta potential between the nanocarriers. Additionally, morphological differences, pertaining to both size (*38*) and shape (*39, 40*), have been implicated in influencing the electrophoretic mobility of a particle in solution. This latter effect may contribute to the differences observed between Phos-MCs and Phos-FMs, as the orientation of high aspect ratio structures, like FMs, in the direction of the electric field can result in an increase in the observed electrophoretic mobility (*39, 40*). Finally, previous studies have shown that the magnitude of the zeta potential can be influenced by particle concentration (*41-44*). The nanocarrier formulations in our studies are controlled on BCP concentration rather than particle concentration, however, we acknowledge that the contribution of nanocarrier concentration can influence the observed differences in zeta potential as well. Relevant physicochemical characteristics for both spherical and cylindrical nanocarriers are presented in **Table S1**.

### Human Plasma Protein Adsorption to PEG-b-PPS Nanocarriers

Nanocarrier characteristics including morphology (*15*), size (*45*), radius of curvature (*46*), surface charge (*18*), and surface chemistry (*47*) have been implicated in governing both the composition and amount of protein adsorbed to the nanocarrier surface following its introduction into a biological system. This new biological identity resulting from the protein corona will eventually dictate the effectiveness of the engineered construct at both reaching (*48, 49*) and interacting with its cellular target (*50*). We first assessed protein adsorption to our nine nanocarrier types and investigated whether their physicochemical properties result in compositional differences in protein coronas formed in human plasma (**Fig. 2A**). Our analysis utilized a combination of traditional protein characterization methods, as well as modern proteomics techniques (**Fig. 2A**). Previous studies demonstrate compositional changes in the PC can result from the choice of surrogate biological milieu (*51*). These compositional changes, which include increased abundances of opsonins like complement proteins and fibrinogen in the nanocarrier PC (*51*), are of consequence for studies conducted using immune cells (*52*). Therefore, we utilized plasma as it best recapitulates nanocarrier molecular interactions *in vivo*.

**Fig. 2.**
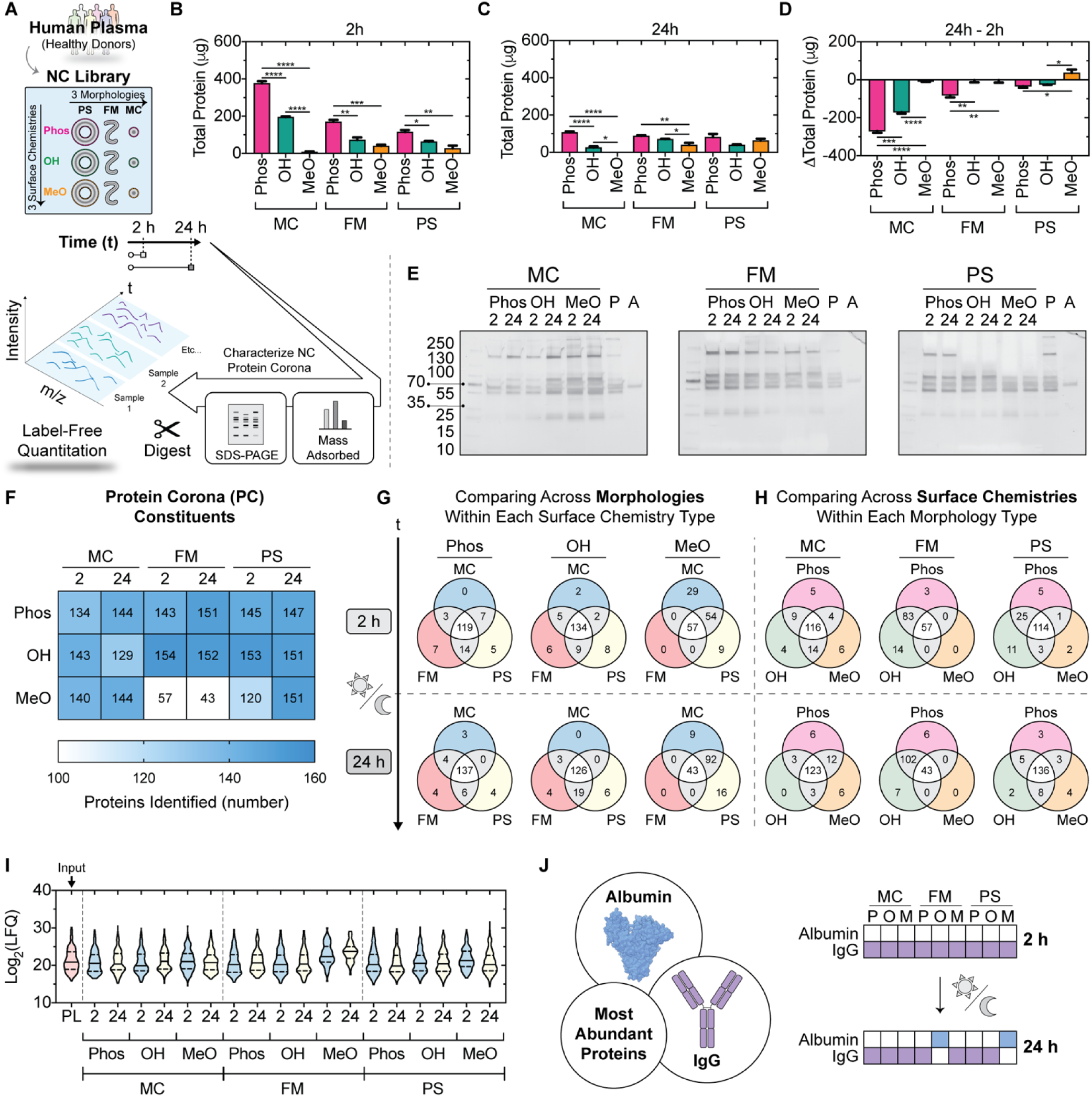
Overview of proteomic analysis of human plasma proteins adsorbed to each soft nanocarrier after 2 h and 24 h. (**B, C**) Total adsorbed protein measured in nanocarrier-protein complexes isolated after incubation with human plasma for two hours (**B**) and 24 hours (**C**). (**D**) Difference in total adsorbed protein at the 24 h timepoint relative to the 2 h timepoint. For all cases, error bars represent s.e.m., n = 3. *p<0.05, **p<0.01, ***p<0.001, and ****p<0.0001. (**E**) Silver stained adsorbed proteins separated by SDS-PAGE grouped by micelle (left gel), filomicelle (middle gel), and polymersome (right gel) morphologies. 1 μg of total protein was run per lane for each sample. (**F**) The total number of each protein corona identified by label-free quantitation. (**G, H**) Venn diagrams depicting common and shared constituents identified in each protein corona. Morphological comparisons within each surface chemistry are displayed in (**G**), whereas (**H**) displays surface chemistry comparisons within each morphology. (**I**) Violin plots depicting the distribution of relative abundances of protein corona constituents. The log2 label free quantitation (LFQ) is displayed as a readout of protein relative abundance. (**J**) The top ranked protein corona constituent by relative abundance at the 2 h and 24 h timepoints.

Nanocarriers (5 mg/mL polymer concentration) were incubated 1:1 with human plasma pooled from over 200 healthy donors to form nanocarrier-protein complexes (see **Fig. S6-S7** for procedure optimization). Significant changes in total adsorbed protein were observed between different surface chemistries within a specified morphology at the 2 h (**Fig. 2B**) and 24 h (**Fig. 2C**) time points. For MC and FM morphologies, the lowest level of protein adsorption was consistently observed for the MeO-functionalized nanocarriers at each timepoint (**Fig. 2B, C**). This suggests that nanocarriers self-assembled from polymer terminated with a methoxy group are most resistant to protein adsorption. A departure from this trend was observed for the vesicular nanocarriers (PS) at 24 h, where the minimum adsorption was found for OH PS, rather than MeO PS (**Fig. 2C**). Interestingly, MeO-functionalized PS were the only morphology-surface chemistry combination that accumulated protein with time, whereas adsorption decreased over time for all other nanocarriers (**Fig. 2D**). Protein adsorption influenced nanocarrier agglomeration, which was specifically observed for the OH-and MeO-functionalized PSs after exposure to human plasma for 2 h and 24 h (**Fig. S8**). For MeO-PS, the increase in adsorbed protein at 24 h coincided with nanocarrier agglomeration (**Fig. S8**). These macroscopic observations suggest surface chemistry-mediated differences in protein adsorption to PEG-*b*-PPS nanocarriers might influence their colloidal stability, which holds consequences on their cellular interactions and immunogenicity.

The overall time-dependent decrease in protein adsorption observed for eight of the nine nanocarrier formulations suggests, in most cases, that a transient spike in adsorption occurs upon nanocarrier introduction into the protein-rich human plasma. Significant decreases in adsorption are observed as equilibrium is established between the free and adsorbed protein subpopulations in the nanocarrier-plasma mixture (**Fig. 2B-D**). The relative change in total adsorbed protein was only significant between all examined surface chemistries of the MC morphology (**Fig. 2D**). For FMs, the loss in protein adsorbed to the anionic surface was significantly greater than that of the neutral surfaces (**Fig. 2D**). In general, neutral surfaces had the most stable level of adsorbed protein between timepoints across all nine nanocarrier formulations examined (**Fig. 2D**).

The effect of protein adsorption on nanocarrier surface charge differed at the two timepoints examined. After 2 h, protein adsorption shifted the nanocarrier surface charge in the negative direction for six of the nine formulations, whereas seven of the nine formulations became more positively charged by the 24 h timepoint (**Table S2**). In other words, short-term (≤ 2 h) protein adsorption increased the magnitude of negative charge for the majority of the nanocarriers examined, while adsorption/desorption events occurring over longer timescales had the opposite effect (**Table S2**). The biomolecules making up the plasma represent the ‘change agents’ capable of driving alterations in the nanocarrier surface properties upon interaction. The majority of plasma proteins have isoelectric points that yield an overall negative charge at physiological pH (*53-56*). This result is therefore consistent with the reduction in total protein observed for the majority of the formulations at the 24 h timepoint (**Fig. 2D**), since the loss of protein (predominantly of negative charge) coincides with a shift in NC zeta potential in the positive direction.

Having quantified bulk protein adsorption and associated physicochemical changes to the nine nanocarrier formulations, we sought to examine whether the bulk composition and relative abundance of adsorbed proteins differed depending on morphology, surface chemistry, and plasma incubation time. To this end, one microgram of total protein from each formulation was separated by SDS-PAGE and proteins were detected by silver-staining (**Fig. 2E**). The PEG-*b*-PPS BCPs are undetectable by silver staining and therefore do not interfere with this analysis (**Fig. S9**). For proteins electrophoresed in one dimension, a single band may correspond to multiple protein species having similar molecular weights that are not resolvable by the polyacrylamide matrix. In any case, this analysis permitted the comparison of big picture differences in the composition of proteins adsorbed to each structure in our NC library. Differences in the number and relative intensity of protein bands were observed across each formulation (**Fig. 2E**). Interestingly, high molecular weight protein species (>180 kDa) were observed in all MC and FM formulations, yet were variable among PS (**Fig. 2E**). In particular, proteins of >200 kDa were not detected in the PS corona by silver staining, regardless of PS surface chemistry (**Fig. 2E**).

Given the limitations of simple gel-based methods in conducting large-scale protein investigations, we used label-free proteomics techniques to identify protein constituents of each corona formed in human plasma, and quantified their relative abundances. This analysis identified a total of 171 proteins using a 1% false discovery rate (**Fig. S10**). As expected, the majority of identified proteins had a molecular weight <100 kDa (**Fig. S11**), consistent with the corresponding distributions observed by SDS-PAGE (**Fig. 2E**). The number of identified proteins ranged between 43-154 protein species, depending on the nanocarrier formulation and timepoint examined (**Fig. 2F**). The MeO FM corona contained the fewest number of adsorbing proteins (57 proteins at 2 h and 43 proteins at 24 h), compared to >100 proteins for the eight other nanocarrier morphology and surface chemistry combinations (**Fig. 2F**). For surface chemistry groupings (Phos, OH, MeO) and morphology groupings (MC, FM, PS), the number of adsorbing species was not significantly different when considered individually as bulk groups (**Fig. S12**).

Trends in the time-dependent protein corona composition differed between surface chemistry (**Fig. 2G**) and morphology (**Fig. 2H**) groupings. Within surface chemistry groups, neutral surfaces enriched a smaller subset of shared proteins, as observed by the decrease in shared constituents for OH and MeO nanocarriers, while the number of shared adsorbing species increased from 119 proteins (2 h) to 137 proteins (24 h) for anionic Phos surfaces (**Fig. 2G**). Within morphology groups, the number of shared constituents increased for spherical nanostructures (MC, PS), whereas a loss in shared constituents was observed for the cylindrical FM morphology (**Fig. 2H**). The overall shape of the relative abundance distributions further demonstrates compositional differences between nanocarrier formulations, and these distributions differed from that of the pooled human plasma input sample (**Fig. 2I**). Protein relative abundance generally did not correlate across formulations (**Fig. S13-S14**). Two blood proteins were found as the most abundant protein corona constituent in all cases: serum albumin (ALB) and immunoglobulin G (IgG) (**Fig. 2J**). IgG dominated at the 2 h timepoint, whereas albumin displaced IgG as the most abundant constituent of the OH FM and MeO PS protein coronas at 24 h (**Fig. 2J**). Aside from these proteins, the serum amyloid P-component (APCS), apolipoprotein A-I (APOA1), complement C1q subcomponent subunit C (C1QC; P02747), and complement C3 (C3; P01024) were also found in high abundance (discussed later in the manuscript). Collectively, these results suggest polar surface chemistries are generally more susceptible to protein adsorption than nanocarrier surfaces terminated with methoxy groups, as this trend was found across all soft spherical and cylindrical nanostructures included in our investigation. The number of adsorbing species, trends in shared versus unique protein corona constituents, and their relative abundances differed depending on the combination of nanocarrier morphology and surface chemistry.

### Characterization of nanocarrier biological identities after 2 h of protein absorption

Considering that the majority of intravenously administered nanocarriers are cleared systemically within a few hours of injection (*17, 33, 57, 58*), the protein corona that forms during this early time period represents a critical biological identity that strongly influences nanocarrier fate and performance. We therefore examined high level gene ontology (GO) terms associated with the set of differentially enriched proteins across all nine nanocarriers at the 2 h absorption timepoint following incubation with human blood (**Fig. 3A, B**). Significant differences in protein corona relative abundances was found for a set of 121 proteins. This set of proteins is significantly associated with biological process terms of stress response, immune system processes, and transport (**Fig. 3B**). A more detailed network analysis reveals significant involvement of these 121 differentially adsorbed proteins in blood coagulation and complement activation biological processes, as well as wounding and inflammatory responses (**Fig. 3C**; see **Fig. S15** for the full node annotations of the network).

**Fig. 3.**
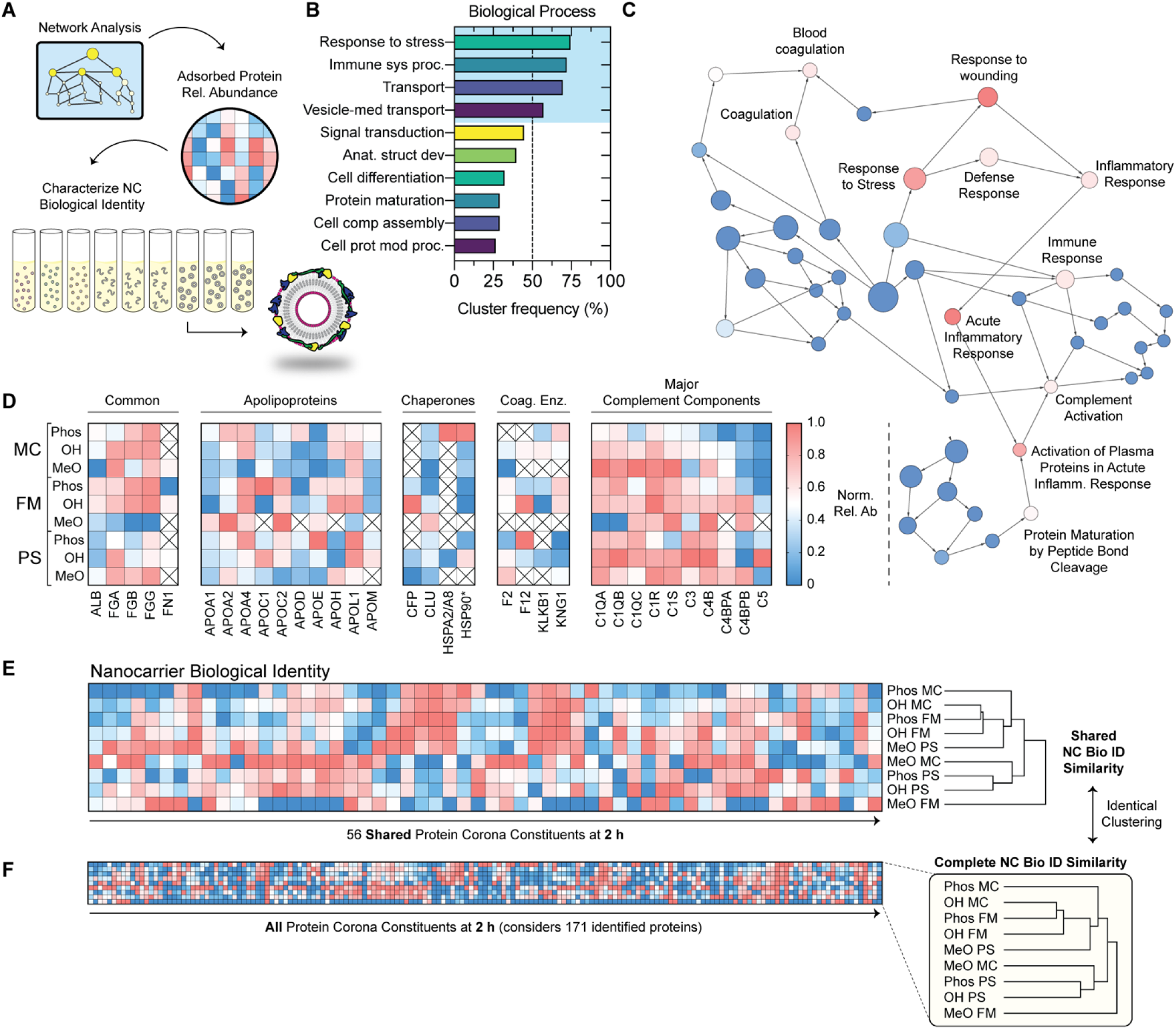
The biological identity of soft polymeric nanocarriers in human plasma at the 2 h timepoint is dependent on the combination of morphology and surface chemistry. (**A**) Illustrative overview. (**B**,**C**) Biological process gene ontology (GO) analysis of the set of 121 differentially enriched proteins across all nine NCs (ANOVA, *q*<0.01). (**B**) The top ten overrepresented GO slim terms (high level nodes). (**C**) Network analysis of overrepresented protein corona constituents. The network was constructed from highly significant terms (corrected p-value < 1e-20) to simplify presentation. The 10 most significant nodes are annotated. (**D**) Relative abundance of protein corona subsets with clear physiological significance. Common constituents, apolipoprotein, chaperone, coagulation enzymes, and major complement components (upstream focus) groups are displayed. *HSP90 refers to HSP90AA1/AB1. (**E**,**F**) Hierarchical clustering of the 56 shared constituents (**E**) detected in all nine coronas, and the set of all 171 constituents (**F**) detected in the proteomic analysis. In both cases, hierarchical clustering (Euclidean distance metric) was performed using the average linkage method to represent cluster proximity.

Compositional differences were found in many different protein groupings, including common adsorbing proteins (serum albumin, fibrinogen, fibronectin), as well as apolipoproteins, protein aggregation and complement-responsive chaperones, coagulation enzymes, and major complement components (**Fig. 3D**). A greater enrichment of fibrinogen was observed compared to albumin at 2 h, and fibronectin was only found in micellar (MC, FM) coronas with the exception of OH PS (**Fig. 3D**).

Interestingly, C1QC, APCS, and C3 were dominant constituents of the respective MeO MC, Phos PS, and OH PS coronas, suggesting strong interactions with these surfaces (**Fig. 3D**). The biological chemistry that underlies the C3 observation is particularly interesting. The polar surfaces (Phos, OH) included in this investigation contain hydroxyl groups capable of nucleophilic attack on C3, whereas the non-polar MeO surface lacks this capability. This observation may suggest the polar nanocarrier surfaces may arise from covalent attachment between C3 and the hydroxyl groups of the OH PS and Phos PS polymers.

Biological identities are complex fingerprints defined by both the constituency and relative abundance of numerous adsorbing plasma proteins. To compare the similarity of protein coronas formed on PEG-*b*-PPS nanocarriers at 2h, we performed hierarchical clustering on two different subsets of proteins: the set of all shared protein corona constituents (**Fig. 3E**) and the inclusive list of all 171 proteins identified by our proteomic investigation (**Fig. 3F**). In this analysis, nanocarrier protein coronas that cluster together are more similar than those that do not.

The dendrogram branching patterns were identical for both cases (**Fig. 3E,F**), suggesting the differential abundance of the 56 proteins common to all nine NCs is sufficient to capture the defining features of each biological identity at 2 h. The major difference was that MeO FM did not cluster with the remaining nanocarrier morphology and surface chemistry combinations (**Fig. 3E,F**). This suggests the biological identity of MeO FM is the most unique, which was suggested by our initial, more crude analysis of protein corona constituency (**Fig. 2F**).

For the remaining eight nanocarrier types, MeO MC clustered with polar vesicular structures, whereas the polar spherical and filamentous micelle structures (Phos MC, OH MC, Phos FM, and OH FM) clustered with the non-polar MeO PS vesicular structures (**Fig. 3E,F**). The former analysis therefore suggests non-polar micelles have a similar biological identity to polar vesicles, whereas the latter analysis suggests the biological identity of polar micellar spheres and cylinders are most similar to that of non-polar vesicles.

From these analyses, we conclude that the biological identity of soft nanocarriers in human blood depends on the combined effect of morphology and surface chemistry on its interactions with plasma proteins. The relative abundance profile of the subset of 56 shared protein corona constituents (**Fig. 3E**) is sufficient to capture the biological identity that includes all quantified proteins (**Fig. 3F**). These biological identities, interpretable as composite molecular signatures unique to each nanocarrier morphology and surface chemistry combination, are the molecular surfaces that interface with cells and tissues.

### Assessment of the Inherent Inflammatory Activity of PEG-*b*-PPS Nanocarriers in Human Blood Samples

To better understand the immunogenicity of soft nanocarriers and their molecular determinants, we examined complement activation and cytokine responses (**Fig. 4A**) induced by each nanocarrier type in human blood samples. We hypothesized that nanocarrier protein corona composition could predict downstream complement activation. Specifically, we investigated whether nanocarriers having similar complement protein compositions at an early 2 h timepoint would generate similar levels of C3a, C4a, and/or C5a peptide cleavage products (anaphylatoxins). We further hypothesized that anaphylatoxin-inducing nanocarriers would elicit significantly higher secretion of pro-inflammatory cytokines from PBMCs. Complement activation was assessed by incubating each of the nine nanocarriers with pooled human sera. Afterwards, the concentration of C3a, C4a, and C5a was determined in collected serum. To our surprise, there were no statistically significant differences in the concentration of C3a for serum incubated with any nanocarrier formulation in comparison to PBS-treated serum (**Fig. 4B**). Previous studies demonstrate both hydroxylated and negatively charged nanocarrier surfaces can activate complement through the alternative pathway (*59, 60*). Polyhydroxylated nanocarrier surfaces are thought to contribute to the activation of the alternative pathway of complement by inducing a nucleophilic attack of the internal thioester bond exhibited by C3b (*61*), which can accelerate the alternative pathway through C3 tick-over (*62*). Considering this, we expected that the OH-functionalized and, potentially, the Phos-functionalized nanocarriers would activate complement via the alternative pathway. Notably, the C3a levels induced by PEG-*b*-PPS nanostructures in these experiments, per polymer concentration, were substantially lower than those elicited by dimyristoyl-phosphocholine (DMPC) liposomes(*63*). While no statistically significant differences were observed for the serum concentration of C3a, significant increases in C4a (**Fig. 4C**) and C5a (**Fig. 4D**) were observed with respect to a PBS-treated control.

**Fig. 4.**
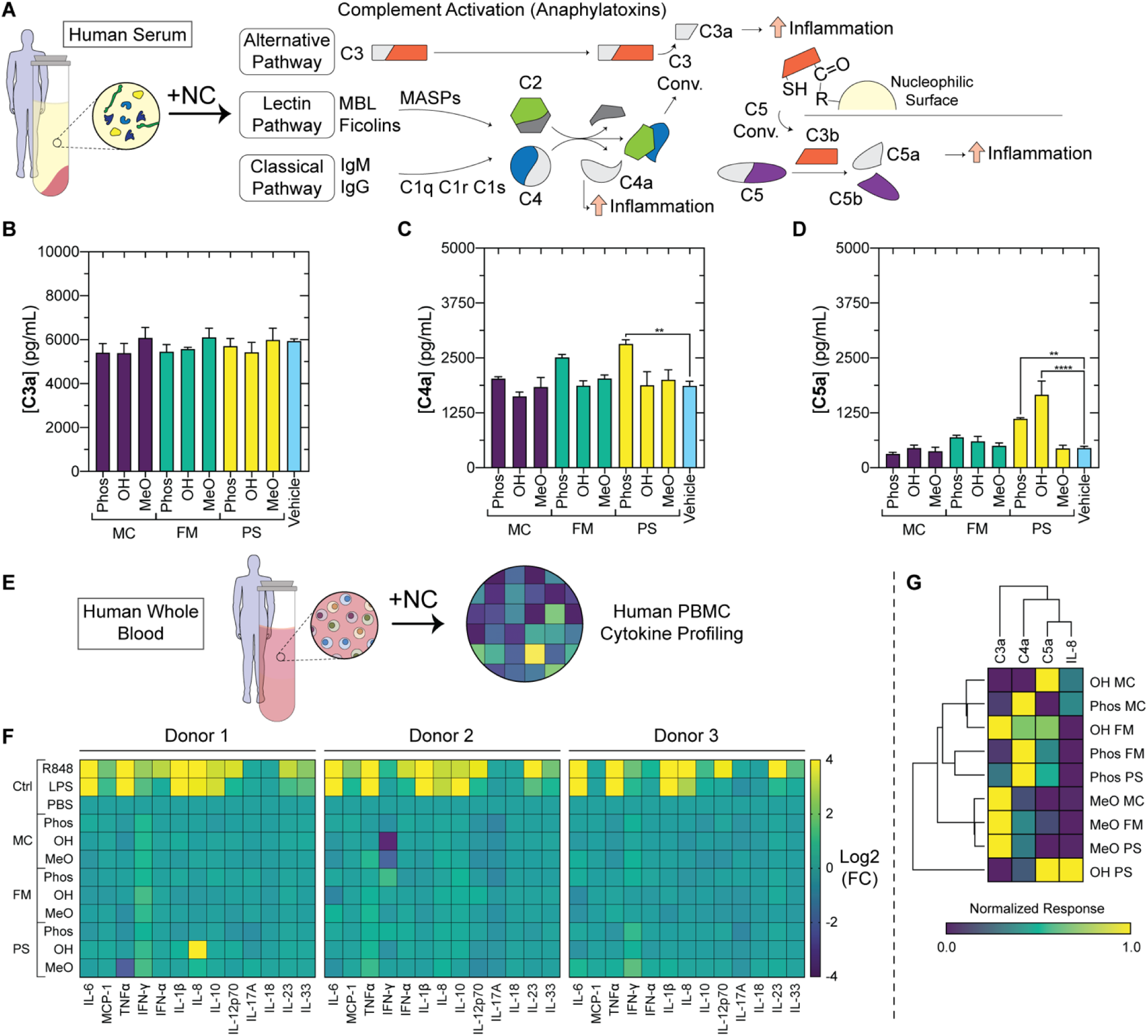
Assessment of nanocarrier complement activation and immunogenicity upon introduction into human blood samples. (**A**) Illustration of complement activation and pro-inflammatory responses. (**B-D**) Assessment of complement activation after nanocarrier introduction into pooled human serum. The serum concentrations of (**B**) C3a, (**C**) C3a, and (**D**) C3a anaphylatoxins were determined (n = 3). Error bars represent s.e.m. Significance was determined with Tukey’s multiple comparison test. \*\**p*<0.01, \*\*\*\**p*<0.0001. (**E**) Illustration of cytokine profiling experiment. Cytokine levels were assessed after incubating nanocarriers in human whole blood samples obtained from individual donors (n = 3). (**F**) Heatmap depicting changes in PBMC cytokine secretion levels induced by nanocarrier incubation with whole blood for 4 h. (**G**) Hierarchical clustering of nanocarriers based on C3a, C4a, C5a, IL-8 responses. Each response was normalized to a 0.0-1.0 scale. Normalization procedures included PBS negative control. Hierarchical clustering was performed using the Euclidean distance metric and average linkage methods.

Phos-functionalized surfaces consistently induced greater increases in C4a concentration than OH and MeO surfaces (**Fig. 4C**). However, the C4a induction was only significant in the case of Phos PS (**Fig. 4C**). Charge is a particularly strong contributing factor for classical pathway activation, but does not completely explain the greater C4a generation by Phos PS, as Phos FM and Phos MC were also highly negatively charged but did not elicit significant responses compared to the PBS control (**Fig. 4B-D**). Previous studies demonstrated anionic lipid vesicles activate the classical pathway through direct binding of C1q (*64*), an opsonin contributing to initiating the classical arm. C1q is composed of six subunits with globular cationic heads that likely use electrostatic interactions to directly interface with anionic surfaces (*64*). This interaction has been proposed for vesicular structures, but studies involving lipid vesicles have shown that nanocarrier-C1q interactions depend on the specific anionic functionality involved rather than simply the nanocarrier surface charge (*65*). Previous studies involving carboxylic acid-functionalized, anionic lipid MCs did not observe the direct interaction of C1q with the MC surface, potentially suggesting that the high curvature of MCs inhibits these structures from interacting with C1q (*66*). Consistent with these findings, our proteomic investigations found collectively higher relative abundances of C1QA, C1QB, C1QC in the protein corona formed on Phos PS (vesicular structures) than Phos MC (**Fig. 3E**). The relative abundance of these C1Q proteins exceeded a normalized value of 0.5 for all three detected proteins (**Fig. 3E**). This observation of a greater response to Phos PS compared to Phos FM and Phos MC suggests a structural dependency on complement activation by nanocarriers.

C5a is the most potent of the three anaphylatoxins and is a downstream product of all three arms of the complement cascade. C5a is involved in both the recruitment and activation of neutrophils (*67*) and is implicated in the amplification of IL-8 secretion. Only Phos-and OH-functionalized PSs induced statistically significant increases in C5a serum concentration (**Fig. 4D**). The C5a concentrations that result after incubation with Phos PS and OH PS correspond to increases of approximately 150% and 270%, respectively (**Fig. 4D**).

Despite differential complement activation, we note that the C3a, C4a, and C5a activation induced by PEG-*b-*PPS was low (<10 ng/mL in all cases; **Fig. 4B-D**). The absolute concentrations reported here are orders of magnitude lower than those reported elsewhere in the literature for Zymosan, an established activator of the complement system, which elicited average C3a, C4a, and C5a concentrations in the range of 150-200, 150-200, and 60-80 ng/mL, respectively (*68*). To assess pro-inflammatory responses, we incubated nanocarriers with whole blood samples from three healthy individuals (**Table S3**; **Fig. S16**) and measured PBMC-secreted cytokine profiles after 4 h (**Fig. 4E, F**). Cytokine levels were also measured at 20 h to assess the persistence and dynamics of the PBMC response to each nanocarrier (**Fig. S16-S17**). After incubation, plasma was collected and analyzed for proinflammatory cytokines, including interleukin (IL)-6, tumor necrosis factor (TNF)-α, interferon (IFN)-γ, IFN-α, IL-1β, IL-12p70, IL-17A, IL-18, and IL-33, proinflammatory chemokines monocyte chemoattractant protein (MCP)-1 and IL-8, and the anti-inflammatory cytokines IL-10 and IL-23. Irrespective of their unique combination of morphology and surface chemistry, PEG-*b*-PPS nanocarriers failed to induce a statistically significant alteration in the levels of secreted cytokines when compared to PBS treatment or lipopolysaccharide (LPS) and R848 positive controls (**Fig. 4F**). Compared to PBS control, OH-functionalized PSs induced substantial increases in IL-8 concentration in donor 1 following plasma incubation times of 4 and 20 hours (**Fig. 4F**; **Fig. S16, S17**). We suspected that the increase in IL-8 secretion resulted from C5a production via the complement cascade. IL-8, also known as C-X-C Motif Chemokine Ligand 8 (CXCL8), is a chemokine secreted by monocytes, polymorphonuclear cells, and endothelial cells that is responsible for neutrophil recruitment during inflammation (*69*). Interestingly, IL-8 secretion in whole blood was greatest in response to Phos PS and OH PS at the 4 h timepoint (**Fig. 4F**). A hierarchical clustering analysis of normalized anaphylatoxin and IL-8 responses suggests a link between the C5a and IL-8 responses and the greater immunogenicity of OH PS observed across multiple independent experiments (**Fig. 4G**).

While these results suggest the complement activating capabilities of Phos-and OH-functionalized PEG-*b*-PPS PSs influence additional inflammatory pathways, further studies are required to verify the interplay between the induced secretion of C5a and IL-8 in this BCP system. Closer inspection of the individual donor cytokine profiles revealed the IL-8 increase was not the result of a consistent uptick in chemokine secretion amongst all donors, but rather the result of a particularly strong response from one individual donor (**Fig. S16**). The general variability among the three donor’s responses, where in some instances converse responses for the same cytokine are observed, highlight the necessity of exploring a personalized approach for the development of patient-specific platforms. However, further development of such an approach extends outside the scope of this work.

Lastly, hierarchical clustering further suggests the combination of cytokine responses evoked by OH PS is unique (**Fig. 4G**). Aside from OH PS, clustering nanocarriers by complement and IL-8 responses revealed similar responses for all nanocarrier chassis possessing MeO surface chemistry despite their differences in morphology (**Fig. 4G**). This clustering also revealed the similarity of large Phos-functionalized FM and PS that induced greater C4a responses (**Fig. 4C**). The immunogenicity of polar MCs was most similar to that of OH FM (**Fig. 4G**). We conclude that the surface chemistry of soft nanocarriers is the strongest physicochemical determinant of nanocarrier complement activation and potentially adverse cytokine responses in our studies. Taken together with our proteomic studies, polar chemical groups with a greater potential for electrostatic interaction with C1q proteins and/or covalent attachment with C3 (**Fig. 3D**) were more susceptible to inducing anaphylatoxins and IL-8 responses. In terms of rational design rules for soft nanocarrier engineering, the choice of morphologies with a higher radius of curvature, such as the micelles, might provide a means for mitigating some C1q-associated immunogenicity concerns if polar anionic surface chemistries are desirable due to application-specific advantages.

### Compositional Dynamics of PEG-*b*-PPS Nanocarrier Protein Coronas

Protein adsorption is a dynamic process. The compositional evolution of the protein corona arises from the continuous association and dissociation of proteins with nanocarrier surfaces as the system equilibrates. Protein adsorption/desorption processes on polymer surfaces occur over tens of hours (*70, 71*). The 2 h and 24 h time points used in our studies therefore capture the compositional differences on short and long-time scales. Hierarchical clustering was performed on the shared (**Fig. 5A**) and complete (**Fig. 5B**) protein corona constituents to examine nanocarrier biological identity at 24 h. The biological identity of MeO FM was the most distinct (**Fig. 5A, B**), which is similar to our observations at 2 h (**Fig. 3E, F**). This uniqueness of MeO FM may be due to the substantially fewer constituents that comprise its protein corona (**Fig. 3F**). The low level of protein adsorption to MeO FM (**Fig. 2B, C**), and its lower compositional diversity compared to other nanocarriers (**Fig. 2F**), suggests the synthetic identity of this nanocarrier morphology and surface chemistry combination is most well preserved in human blood compared to the other formulations examined. It is notable that the MeO PS was also found to be relatively unique (**Fig. 5A, B**). Considered together with the MeO FM observation, these results suggest larger nanostructures (PS, FM) with non-polar surfaces develop more unique biological identities at later timepoints. Neither of these structures elicited significant complement activation and cytokine responses in human blood (**Fig. 4**), suggesting the unique protein features of these coronas do not elicit significant negative molecular and cellular responses.

**Fig 5.**
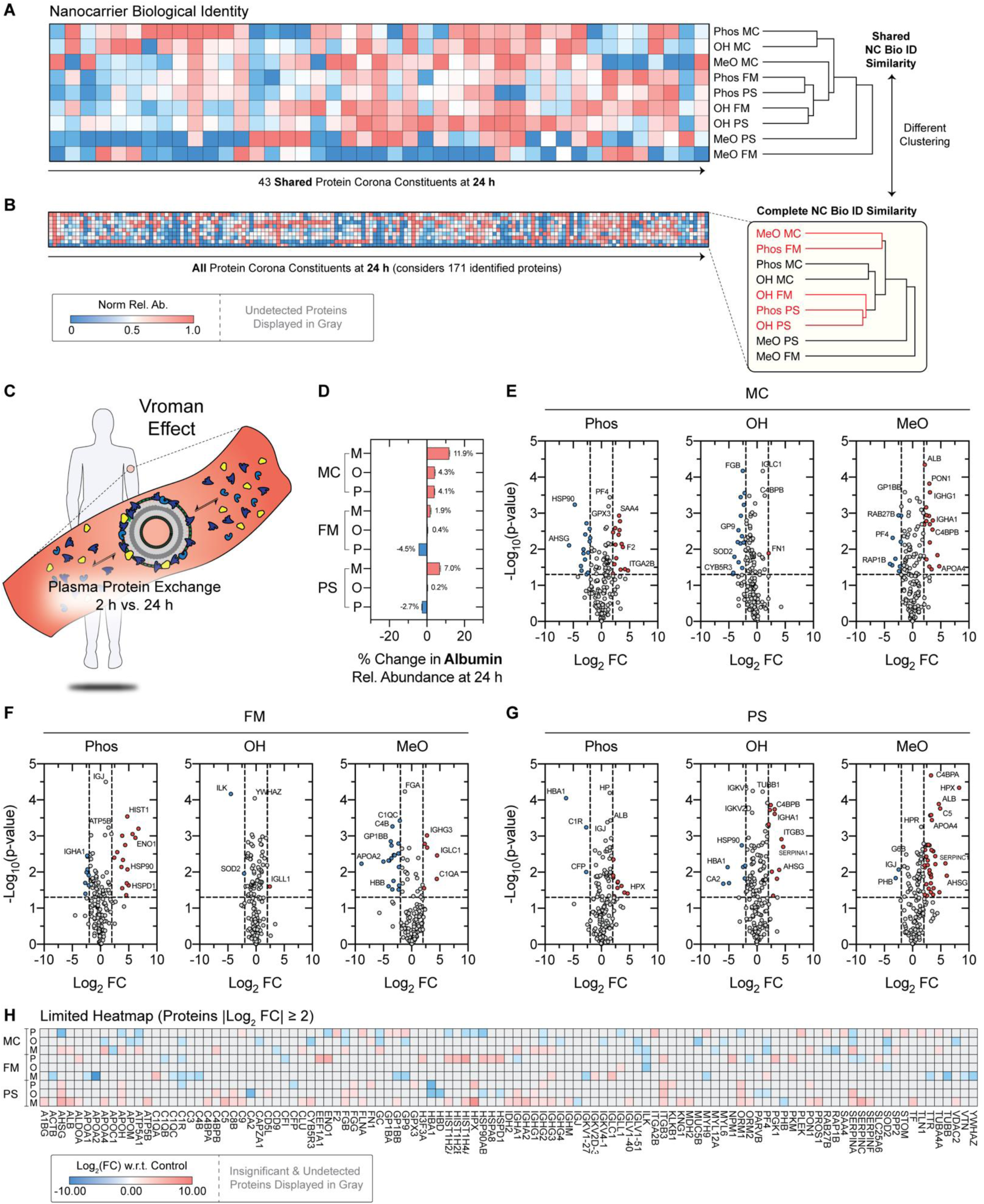
Analysis of time-dependent dynamics in the composition of soft nanocarrier protein coronas. (**A, B**) Hierarchical clustering of nanocarrier biological identity defined by the (**A**) shared 43 proteins adsorbed to all nine nanocarriers at 24 h, and (**B**) the complete set of protein corona constituents quantified at 24 h. (**C**) Vroman effect illustration. (**D**) Percent change in albumin relative abundance at 24 h. (**E-G**) Volcano plots displaying proteins with significant changes in relative abundance at 24 h. The -log10 of the *p*-value is plotted against the log2 fold change (FC). Proteins with a significant ≥ 2-fold change (FC) in relative abundance at 24 h (compared to 4 h) are displayed in blue (≥ 2-fold decrease in relative abundance) and red (≥ 2-fold increase in relative abundance). The abbreviations of the genes that encode the quantified proteins are displayed as space permits. Gene names are used in place of protein accession numbers for ease of interpretation. Significance was determined by t-test (\**p*<0.05). (**H**) Limited heat map displaying the set of proteins with significantly altered relative abundances. The log2(FC) is displayed.

MeO FM and MeO PS aside, the shared protein corona of polar micelles clustered together at this later 24 h timepoint, as did FM and PS of identical surface chemistry (**Fig. 5A**). In contrast to our 2 h biological identity analysis (**Fig. 3E, F**), differences in dendrograms were observed for the shared and complete biological identities at 24 h (**Fig. 5A, B**). When considering the presence of proteins that were unique to one or more nanocarriers (**Fig. 5B**), the corona formed on the non-polar MeO MC surface became more similar to that of Phos FM (**Fig. 5B**). This suggests that the unique proteins associated with the Phos FM corona may contribute to a divergence in its biological identity from those of polar PS and the OH FM (**Fig. 5B**). Furthermore, the complete OH FM became more similar to that of polar PS (**Fig. 5B**). The complete polar PS biological identities were similar (**Fig. 5B**), and the inclusion of less common constituents differentiated these nanostructures from FM counterparts of identical surface chemistry (**Fig. 5A**).

Next, we sought to investigate the exchange of protein corona constituents at 2 h vs 24 h in greater detail. The “Vroman effect” refers to the exchange of absorbed protein constituents that occurs with time, due to the replacement of early-adsorbing higher concentration proteins with proteins having a greater affinity for the nanocarrier surface. At 24 h, substantial changes in protein corona constituency were observed overall. The relative abundance of albumin increased with time (**Fig. 5D**). Statistically significant changes in the relative abundance of proteins between the two timepoints were determined. Protein corona constituents with greater than or equal to a two-fold change in relative abundance at 24 h are annotated (**Fig. 5E-G**), and the relative abundance differences are displayed in a limited heat map for comparison (**Fig. 5H**). Interestingly, MeO PS experienced the largest gain in protein corona constituency (**Fig. 5G**), consistent with the time-dependent gain in protein adsorption (**Fig. 2B-D**) and may contribute to the agglomeration (**Fig. S8**) observed for this formulation through promoting protein denaturation and/or aggregation. Collectively, these analyses demonstrate that trends in biological identity at 24 h differ from those at 2 h for soft polymeric nanocarriers. Consistent with our 2 h analysis (**Fig. 3**), we conclude that the biological identity depends on the combination of nanocarrier morphology and surface chemistry at 24 h as well.

### Uptake of PEG-*b*-PPS Nanocarriers by Human MPS Cells

Modulating the structure of proteins adsorbed to soft nanocarriers has been established as one mechanism for modifying soft nanocarrier biodistribution and clearance by the MPS (*17*), yet leveraging steady-state compositional differences for this purpose has not been explored in detail. We therefore sought to determine whether combinations of morphology and surface chemistry could generate compositional differences in nanocarrier biological identity (**Fig. 2-3, 5**) that would achieve differential uptake of PEG-*b*-PPS nanocarriers within human monocytes, macrophages, and dendritic cells. We first assessed the cytotoxicity of each nanocarrier formulation. Monocyte-like THP-1 cells were incubated with nanocarriers at polymer concentrations ranging from 0.1 to 1.0 mg/mL for 24 h. Irrespective of the nanocarrier morphology and surface chemistry combination, the viability of treated THP-1 cells exceeded 80% for all material concentrations tested (**Fig. S18**). For all nanocarriers, the concentration dependent differences in cell viability were not significant within the examined range (**Fig. S18**). The non-cytotoxic nature of PEG-*b*-PPS nanocarriers observed here is consistent with previously published studies (*22, 23, 33, 72*). To ensure that cytotoxicity would not influence our ensuing cell uptake studies, we used a BCP concentration of 0.5 mg/mL. Cell viability exceeded 85% for all formulations administered at this concentration (**Fig. S18**).

We assessed how changes to the protein corona influence nanocarrier interactions with monocyte-like THP-1 cells, macrophage-like differentiated THP-1 cells (denoted MΦ), and monocyte derived immature dendritic cells (DC) (**Fig. 6A)**. To be consistent with our proteomic investigations (**Fig. 3-5**), we utilized human plasma to first form protein corona on PEG-*b*-PPS nanocarrier surfaces. Uptake studies then proceeded using serum-free media to focus only on uptake differences mediated by adsorbed human plasma proteins, without confounding effects from other protein sources that are commonly added to the media (for example, fetal bovine serum). Each formulation was administered under three separate conditions: (i) no protein exposure (i.e. pristine nanocarriers) or after a (ii) 2 h or (iii) 24 h pre-incubation with human plasma. Previous studies involving PEG-*b*-PPS nanocarriers demonstrate circulation times on the scale of days can be achieved following intravenous administration (*33*). To assess nanocarrier uptake via flow cytometry, nanocarriers were prepared to incorporate the hydrophobic dye DiI, which is stably retained within PEG-*b*-PPS nanocarriers (*30*). The percentage of cells that internalized nanocarriers (%NC+ cells) was determined after a 4 h incubation period (**Fig. 6, B-D**), and the fold change in uptake mediated by the protein corona was quantified (**Fig. 6E**).

**Fig. 6.**
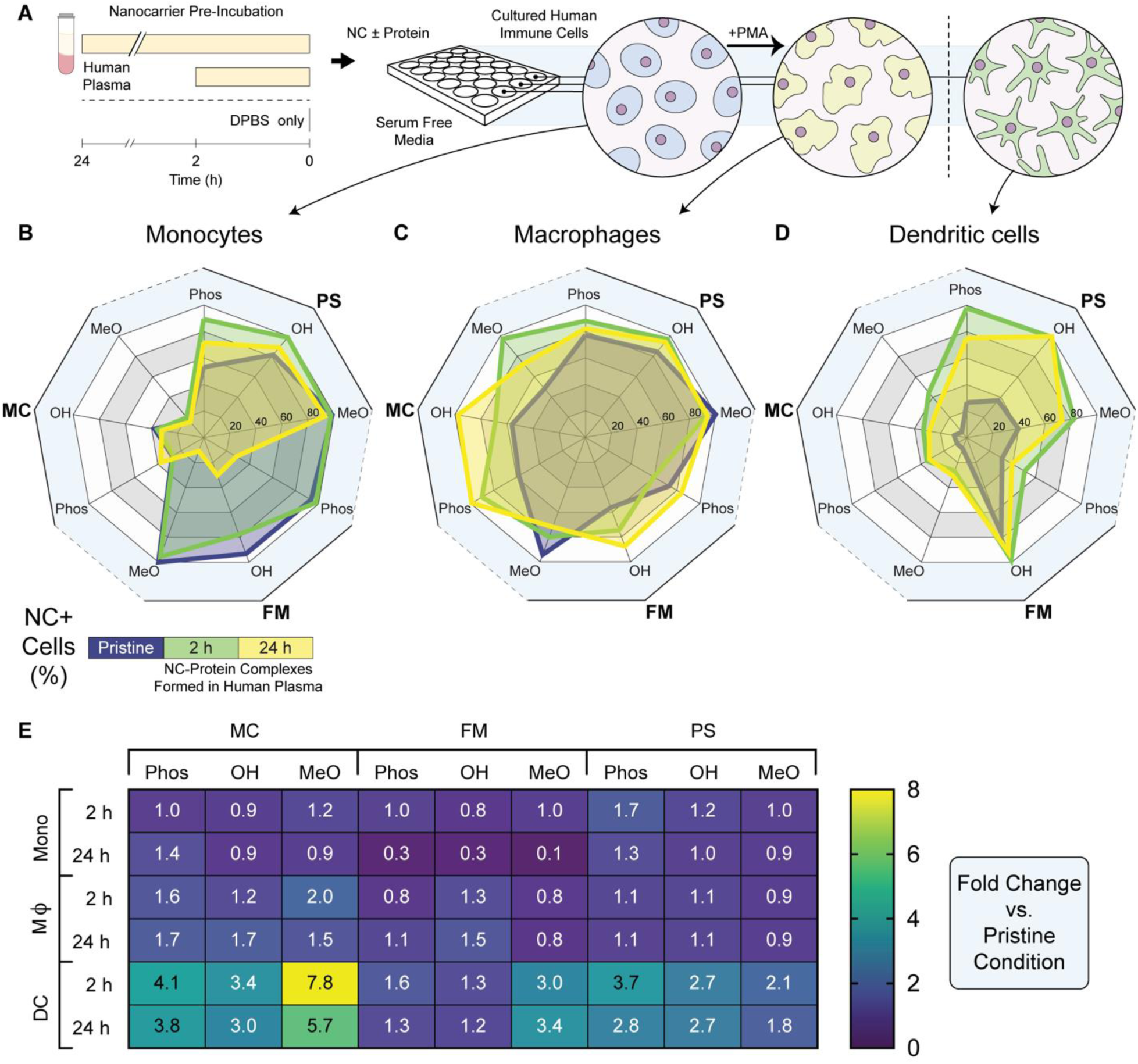
Cellular uptake of nanocarriers in the absence or presence of plasma proteins. (**A**) Schematic depicting process flow for nanocarrier uptake studies. Flow cytometric analysis of nanocarrier (DiI) uptake within (**B**) THP-1 monocytes (Mono; n = 10; except OH-PSs with 24 h plasma incubation where n = 9), (**C**) THP-1 differentiated macrophages (MΦ; n = 10), and (**D**) immature dendritic cells (DC; n = 4). All radar plots depict the percentage of nanocarrier positive cells (% NC+). (**E**) Fold change in % NC+ cells after human plasma protein adsorption. The fold change was calculated w.r.t. the uptake of pristine nanocarriers.

Nanocarrier uptake was first assessed in THP-1 monocytes cells (*73*), which are extensively used to investigate nanocarrier uptake (*50, 74-77*). On the basis of morphology, MCs generally showed significantly less uptake by monocytes than either PSs or FMs (**Fig. 6B**; **Fig. S19A**). This trend was observed when comparing MCs to similarly functionalized PSs under all conditions tested and to FMs under the pristine and 2 h plasma incubation conditions. The protein corona formed at 2 h and 24 h enhanced monocyte uptake of polar PS surface chemistries although changes were much more subtle for MeO PS (**Fig. 6B, E; Fig. S19A**). For FMs, the protein corona formed at 2 h did alter monocyte uptake of cylindrical soft structures, although the magnitude of this effect varied (**Fig. 6B, E**). In contrast, the protein corona formed on FMs at 24 h resulted in a large decrease in uptake by monocytes with respect to the pristine condition (**Fig. 6B, E**). Previous studies investigating the endocytosis of metallic and solid core nanoparticles in THP-1 cells have similarly observed decreases in nanocarrier internalization when serum proteins are present (*50, 74, 75*). These studies suggested that the decreases in uptake were the result of decreases in nanocarrier zeta potential (more negative) upon protein adsorption. Here, statistically significant correlations were not found between soft nanocarrier uptake and zeta potential for any of the condition groupings (**Fig. S20**).

Collectively, the results from our monocyte uptake studies suggest a few trends. Monocyte uptake of micelles is largely unresponsive to the plasma protein coatings irrespective of the timepoint examined within a 24 h period. The protein corona that evolves with time on polar vesicle surfaces renders these structures more prone to monocyte uptake with time, whereas the MeO PS surface develops a protein corona that has small yet significant effects on monocyte uptake. Lastly, the FM morphology develops a protein corona at 2 h with significant effects on monocyte-mediated clearance of nanostructures, whereas the corona formed on these soft cylinders at 24 h decreases monocyte uptake regardless of surface chemistry. These decreases in uptake do not appear to be charge related, suggesting these decreases in FM uptake by monocytes arise due to unique features of the FM biological identity.

Next, we investigated nanocarrier uptake by macrophage-like differentiated THP-1 cells, which are preferable to peripheral blood mononuclear cell (PBMC)-derived macrophages on the basis of their phagocytic capacity (*78*). As such, dTHP-1 cells have been widely used for investigating nanocarrier uptake *in vitro* (*50, 74, 79*). As expected, dTHP-1 cells showed an overall increase in affinity for PEG-*b*-PPS nanocarriers in comparison to their monocyte-like counterparts (**Fig. 6C**). General trends in nanocarrier association and uptake varied based on nanocarrier shape. In PSs and MCs, a net increase in nanocarrier uptake with respect to the pristine condition was observed for all spherical nanocarriers except for MeO-functionalized PSs upon plasma exposure (**Fig. 6C, E**). Changes in PS uptake, irrespective of surface chemistry, were not significantly different on the basis of nanocarrier condition, whereas the protein-mediated increase in MC internalization after plasma exposure was significant (**Fig. 6C, E**; **Fig. S19B**). This finding is consistent with our past observation of high uptake of both MeO MC and MeO PS by macrophages (*33*). The overall increase in spherical nanocarrier uptake by dTHP-1 cells in comparison to their monocyte-like counterparts is most likely due to differences in both scavenger and complement receptor expression (*80*). Cylindrical FMs displayed no major trends in uptake following plasma exposure. On the basis of surface chemistry, plasma exposure led to a net increase in macrophage uptake of all Phos-and OH-functionalized nanocarriers (except Phos-functionalized FMs incubated with plasma for 2 h). Aside from the MCs, plasma exposure decreased macrophage uptake of MeO nanocarriers (**Fig. 6C, E**).

Nanocarrier uptake studies performed using monocyte-derived immature dendritic cells revealed the most interesting trends (**Fig. 6D**). Their phagocytic capacity, ability to efficiently process and present antigens, and ability to initiate T cell immunity have made dendritic cells a desirable cellular target for biomaterial-based delivery systems designed for vaccination (*81*). Under the pristine condition, PEG-*b*-PPS nanocarriers were taken up to a lesser extent by dendritic cells compared to monocyte-like THP-1 or macrophage-like dTHP-1 cells (**Fig. 6, B-D**). Generally, nanocarrier internalization and association decreased with increasing BCP f_PEG_ (i.e. PSs exhibited increased uptake in comparison to FMs, which exhibited increased uptake compared to MCs for a given surface chemistry) (**Fig. 6D**). All nanocarriers, irrespective of the morphology and surface chemistry combination, saw an increase in dendritic cell uptake over the pristine condition after incubation with plasma (**Fig. 6D, E**). These increases in nanocarrier uptake were generally only significant for the PS morphology (**Fig. S19C**). This result is highly significant in the context of our laboratories past work examining MPS uptake preferences of MeO MC, MeO PS, and MeO FM nanostructures (*33*). Those studies demonstrated that the PS morphology has a greater specificity for targeting the dendritic cells *in vivo* compared to MC or FM morphologies (*33*), but the mechanism for this effect was not clear. Here, our uptake studies in the presence and absence of protein-free conditions demonstrate the protein corona formed on the surface of PS nanostructures is the source of this targeting specificity. While differences in nanocarrier uptake by dendritic cells were observed within a given morphology and condition on the basis of surface chemistry, these differences were also generally not significant (**Fig. S19C**).

Interestingly, the cell uptake response to the nanocarrier condition was conserved regardless of cell type (**Fig. 6**). Rather than looking at each condition individually, the combination of the pristine, 2 h plasma incubation, and 24 h plasma incubation conditions can be viewed as the nanocarrier’s conditional uptake profile. When looking at the conditional uptake profiles of the nanocarriers, the most common trend observed involved an increase in uptake from the pristine condition to the 2 h plasma incubation condition followed by a decrease in uptake from the 2 h to 24 h plasma incubation condition. This trend was observed in approximately 60% of the conditional uptake profiles investigated. This result suggests that changes in the protein corona influence how PEG-*b*-PPS nanocarriers interact with cells of the MPS over time.

## Discussion

We conducted a systems level investigation of the relationship between protein adsorption and the biological performance of soft nanocarriers in human-derived samples. Using nine model soft nanocarriers and modern proteomic techniques, we demonstrate the combination of morphology and surface chemistry defines the biological identity of soft nanocarriers in human blood. These differences had consequences for nanocarrier performance, including differential immunogenicity and uptake by human MPS cells. Polar surface chemistries accumulated larger amounts of adsorbed protein and generally enriched complement proteins to a greater extent than nanocarriers with methoxy surfaces. Phosphate-and hydroxyl-functionalized polymersomes led to significant increases in serum anaphylatoxin concentrations, whereas only the hydroxyl-functionalized polymersomes elicited a pro-inflammatory response by PBMCs in human blood. In cellular uptake studies, nanocarrier physicochemical property combinations influenced baseline uptake by monocytes, macrophages, and dendritic cells of the MPS. Redefining the synthetic nanocarrier surfaces with biological identities formed after 2 h or 24 h in plasma resulted in a broad range of decreases and increases in cellular uptake by MPS cell types.

Our results have significant implications for the design of nanocarrier drug delivery vehicles for applications in personalized nanomedicine and targeted immunomodulation. The rational design of synthetic nanocarriers with a well-defined biological identity provides greater control over drug biodistribution, adverse effects, and clinical outcome. Our immunogenicity studies demonstrate that understanding nanocarrier biological identity differences is reasonably predictive of the susceptibility for anaphylatoxin induction. However, these trends, established in pooled human blood samples, did not always translate to pro-inflammatory responses in whole blood from individual patients. This observation underscores the need to conduct even more comprehensive investigations into nanocarrier surface properties, biological identity, and differential immunogenicity in a large number of blood samples from individual patients.

## Materials and Methods

### Chemicals

All chemical reagents were purchased from Sigma-Aldrich St. Louis, MO, USA, unless stated otherwise. Fluorescent antibodies, Zombie Aqua fixable cell viability kit, IC cell fixation buffer, and LEGENDplex™ Human Inflammation Panel were acquired from BioLegend.

### Synthesis of Bn-PPS-*b*-PEG_45_-OMe

Linear poly(ethylene glycol) methyl ether (mPEG_45_ MW 2000) was dried via azeotropic distillation (mPEG_45_ = 40 g, 20.0 mmol, 1 equivalent (EQ)) in toluene using a Dean-Stark trap. The dried solution was removed from heat and allowed to cool to room temperature under vacuum prior to purging the reaction vessel with nitrogen (N_2_). Triethylamine (16.72 mL, 120 mmol, 6 EQ) was added to the stirred solution. After cooling to 0°C, methanesulfonyl chloride (17.74 mL, 100 mmol, 5 EQ) diluted in toluene (80 mL) was added to the stirred solution drop-wise. The reaction was stirred overnight at room temperature under N_2_. Salt produced during the reaction was removed via vacuum filtration of the reaction mixture over a celite filter cake. Toluene was removed through rotary evaporation and the product was precipitated in ice-cold diethyl ether. The precipitate was recovered via filtration, washed with cold diethyl ether, and dried under vacuum. The resulting mesylate-functionalized mPEG_45_ (mPEG _45_-OMs) was stored under N_2_ until further use. ^1^H-NMR (400 MHz, CDCl_3_): δ 4.38 – 4.34 (m, 2H,-C<u>H</u>_2_-O-SO_2_), 3.65-3.60 (s, 180H, PEG backbone), 3.37 – 3.35 (s, 3H, O-C<u>H</u>_3_), 3.07 – 3.05 (s, 3H, SO_2_-C<u>H</u>_3_).

mPEG_45_-*b*-PPS was synthesized for FMs and MCs using a small molecule sulfhydryl initiator. Benzyl mercaptan (For FMs: 53.7 μl, 0.45 mmol, 1 EQ; For MCs: 98.7 μl, 0.83 mmol, 1 EQ), dissolved in anhydrous dimethylformamide, was deprotonated through the addition of sodium methoxide (0.5 M solution in methanol; For FMs: 0.50 mmol, 1.1 EQ; For MCs: 0.92 mmol, 1.1 Eq) to generate a thiolate anion. This thiolate anion was used to initiate the living anionic ring-opening polymerization of propylene sulfide (For FMs: 1.50 mL, 19.94 mmol, 44 EQ; For MCs: 1.50 mL, 18.37 mmol, 22 EQ). The terminal thiolate was endcapped through the addition of mPEG_45_-mesylate (For FMs: 1.57 g, 0.68 mmol, 1.5 EQ; For MCs: 2.00 g, 0.87 mmol, 1.05 EQ) and subsequently stirred overnight. Following the removal of dimethylformamide through rotary evaporation, the product was recovered through methanol precipitation and cooled as needed. Resulting precipitate was recovered and dried under vacuum. ^1^H-NMR (400 MHz, CDCl_3_): δ 7.36 - 7.32 (d, 4H, Ar<u>H</u>), 3.68-3.63 (s, 180H, PEG backbone), 3.41 – 3.39 (s, 3H, O-C<u>H</u>_3_), 3.00 - 2.85 (m, 89H, C<u>H</u>_2_), 2.71 - 2.59 (m, 44H, C<u>H</u>), 1.44 - 1.37 (d, 132H, C<u>H</u>_3_).

### Synthesis of Bn-PPS-*b*-PEG_17_-OMe

Linear poly(ethylene glycol) methyl ether (mPEG_17_, MW 750) was dried via azeotropic distillation (20 g, 26.67 mmol, 1 EQ) in toluene using a Dean-Stark trap. The dried solution was removed from heat and allowed to cool to room temperature under vacuum prior to purging the reaction vessel with N_2_. Triethylamine (18.58 mL, 133.35 mmol, 5 EQ) was added to the stirred solution. After cooling to 0°C, methanesulfonyl chloride (10.32 mL, 133.35 mmol, 5 EQ) diluted in toluene (100 mL) was added to the stirred solution drop-wise. The reaction was stirred overnight at room temperature under N_2_. Salt produced during the reaction was removed via vacuum filtration of the reaction mixture over a celite filter cake. Toluene was removed through rotary evaporation and the product was precipitated in ice cold diethyl ether. The precipitate was recovered via filtration, washed with cold diethyl ether, and dried under vacuum. The resulting mesylate-functionalized mPEG_17_ (mPEG_17_-OMs) was stored under N_2_ until further use.

Thioacetate-functionalized mPEG_17_ (mPEG_17_-TAA) was prepared from mPEG_17_-OMs. In brief, mPEG_17_-OMs (5.0 g, 6.67 mmol, 1 EQ) was dissolved in anhydrous dimethylformamide within an RBF that had been previously evacuated and purged with N_2_. Potassium carbonate (3.69 g, 26.68 mmol, 4 EQ) and thioacetic acid (1.90 mL. 26.68 mmol, 4 EQ) were added sequentially. The reaction was stirred overnight at room temperature. Salt produced during the reaction was removed via vacuum filtration of the reaction mixture over a celite filter cake. Dimethylformamide was removed through rotary evaporation. The product was dissolved in tetrahydrofuran and run on an aluminum oxide column. The collected product was concentrated by removing tetrahydrofuran via rotary evaporation and precipitated in ice cold diethyl ether. The precipitate was recovered via filtration, washed with cold diethyl ether, and dried under vacuum. The resulting mPEG_17_-TAA was stored under N_2_ until further use.

mPEG_17_-*b*-PPS was synthesized for the preparation of PSs via the stable macroinitiator mPEG_17_-TAA. mPEG_17_-TAA (0.50 g, 0.40 mmol, 1 EQ), deprotonated through the addition of sodium methoxide (0.5 M solution in methanol, 0.44 mmol, 1.1 EQ), was used to initiate the living anionic ring-opening polymerization of propylene sulfide (1.00 mL, 12.25 mmol, 30 EQ). The terminal thiolate was endcapped through the addition of benzyl bromide (1.44 mL, 2.0 mmol, 5 EQ) and subsequently stirred overnight. Following the removal of dimethylformamide through rotary evaporation, the product was recovered through methanol precipitation and cooled as needed. Resulting precipitate was recovered and dried under vacuum.

### Synthesis of Bn-PPS-*b*-PEG-OH

α-tosyl-ω-hydroxyl PEG_45_ and PEG_23_ was prepared by adapting a previously published protocol (*82*). PEG (PEG_45_: 30 g, 15 mmol, 1 EQ; PEG_23_: 25 g, 25 mmol, 1 EQ) was dried via azeotropic distillation in toluene using a Dean-Stark trap. Following complete removal of toluene, the dry PEG was flushed with N_2_ and dissolved in anhydrous dichloromethane before being placed on ice. Silver (I) oxide (For PEG_45_: 5.21 g, 22.5 mmol, 1.5 EQ; For PEG_23_: 8.69 g, 37.5 mmol, 1.5 EQ), potassium iodide (For PEG_45_: 1.79 g, 10.8 mmol, 0.72 EQ; For PEG_23_: 2.99 g, 18.0 mmol, 0.72 EQ), and *p*-Toluenesulfonyl chloride (For PEG_45_: 3.00 g, 15.75 mmol, 1.05 EQ; For PEG_23_: 5.00 g, 26.25 mmol, 1.05 EQ) were sequentially added to the vigorously stirring solution. After two hours, the sample was removed from its ice bath and was further allowed to react overnight at room temperature under N_2_. Vacuum filtration over a celite filter cake removed generated salts and silver (I) oxide from the crude product. Dichloromethane was removed through rotary evaporation and the product was precipitated in ice cold diethyl ether. The precipitate was recovered via filtration, washed with cold diethyl ether, and dried under vacuum. The resulting tosylate-functionalized PEG (HO-PEG_45_-OTs) was stored under N_2_ until further use. ^1^H-NMR (400 MHz, DMSO): δ 7.78 - 7.72 (d, 2H), 7.47 - 7.43 (d, 2H), 4.55 - 4.49 (t, 1H, O<u>H</u>), 4.09 - 4.06 (t, 2H, C<u>H</u>_2_-SO_2_), 3.49 - 3.46 (s, 180H, PEG backbone), 2.40 - 2.38 (s, 3H, C<u>H</u>_3_).

Benzyl mercaptan (For FMs: 53.7 μl, 0.61 mmol, 1 EQ; For MCs: 62.7 μl, 0.53 mmol, 1 EQ), dissolved in anhydrous dimethylformamide and deprotonated through the addition of sodium methoxide (0.5 M solution in methanol; For FMs: 0.67 mmol, 1.1 EQ; For MCs: 0.58 mmol, 1.1 EQ), was used to initiate the living anionic ring-opening polymerization of propylene sulfide (For FMs: 2.00 mL, 26.58 mmol, 44 EQ; For MCs: 1.00 mL, 12.25 mmol, 23 EQ). The terminal thiolate was endcapped through the addition of HO-PEG_45_-OTs (For FMs: 4.08 g, 1.69 mmol, 2.75 EQ; For MCs: 3.52 g, 1.45 mmol, 2.75 EQ) and subsequently stirred overnight. Following the removal of dimethylformamide through rotary evaporation, the product was recovered through precipitation in methanol and cooled as needed. Recovered precipitate was dried under vacuum. ^1^H-NMR (400 MHz, CDCl_3_): δ 7.34 - 7.30 (d, 4H, Ar<u>H</u>), 3.65 - 3.60 (s, 186H, PEG backbone), 3.00 - 2.85 (m, 89H, C<u>H</u>_2_), 2.69 - 2.57 (m, 44H, C<u>H</u>), 1.42 - 1.35 (d, 130H, C<u>H</u>_3_).

Benzyl mercaptan (42.0 μl, 0.35 mmol, 1 EQ), dissolved in anhydrous dimethylformamide and deprotonated through the addition of sodium methoxide (0.5 M solution in methanol; 0.39 mmol, 1.1 EQ), was used to initiate the living anionic ring-opening polymerization of propylene sulfide (1.00 mL, 12.25 mmol, 35 EQ). The terminal thiolate was endcapped through the addition of HO-PEG_23_-OTs (3.47 g, 2.65 mmol, 7.5 EQ) and subsequently stirred overnight. Following the removal of dimethylformamide through rotary evaporation, the product was recovered through precipitation in methanol and cooled as needed. Recovered precipitate was dried under vacuum. ^1^H-NMR (400 MHz, CDCl_3_): δ 7.32 - 7.29 (d, 4H, Ar<u>H</u>), 3.64 - 3.60 (s, 86H, PEG backbone), 2.96 - 2.81 (m, 80H, C<u>H</u>_2_), 2.66 - 2.54 (m, 40H, C<u>H</u>), 1.40 - 1.33 (d, 119H, C<u>H</u>_3_).

### Synthesis of PPS-*b*-PEG-Phos BCPs

Lyophilized PPS-*b*-PEG-OH (For PSs: 1.0 g, 0.29 mmol, 1 EQ; For FMs: 1.0 g, 0.19 mmol, 1 EQ; For MCs: 1.0 g, 0.28 mmol, 1 EQ) was massed into a previously evacuated and N_2_ purged reaction vessel. The polymer was dissolved in 10 mL of anhydrous tetrahydrofuran and flushed with N_2_ for one hour. After purging with N_2_, the BCP solution was cooled to 0°C. Phosphorous (V) oxychloride (For PSs: 34.3 μl, 0.37 mmol, 1.27 EQ; For FMs: 22.5 μl, 0.24 mmol, 1.27 EQ; For MCs: 33.3 μl, 0.36 mmol, 1.27 EQ) was added dropwise to the vigorously stirred solution. Following the addition, the solution was removed from its ice bath and further allowed to react overnight at room temperature under N_2_. The reaction was quenched through the addition of 5.0 mL of Milli-Q water and allowed to stir for five minutes. Subsequently, the product was extracted with dichloromethane, dried over sodium sulfate, and filtered. Dichloromethane was removed via rotary evaporation. The resulting product was recovered by precipitation and dried under vacuum overnight. ^1^H-NMR (400 MHz, CDCl_3_): δ 7.24 - 7.22 (d, 4H, Ar<u>H</u>), 3.58 - 3.54 (s, 71H, PEG backbone), 2.88 - 2.73 (m, 79H, C<u>H</u>_2_), 2.59 - 2.47 (m, 40H, C<u>H</u>), 1.33 - 1.26 (d, 117H, C<u>H</u>_3_). ^31^P-NMR (162 MHz, CDCl_3_): δ 0.2 ppm.

### Preparation of FMs and MCs via Thin-Film Hydration

FMs and MCs were generated via thin-film hydration. BCP (either Bn-PPS-*b*-PEG-OMe, Bn-PPS-*b*-PEG-OH, or Bn-PPS-*b*-PEG-Phos) was dissolved in ∼2 mL of dichloromethane (0.5 w/v%) within 2.0 mL clear glass vials (ThermoFisher Scientific). BCP films were formed on the inner surface of the vials by removing the dichloromethane under vacuum. The thin films were hydrated with 1 mL of Dulbecco’s phosphate-buffered saline (1x PBS) (ThermoFisher Scientific). FM assemblies were achieved through gentle agitation overnight at room temperature using a Stuart SB3 rotator. MC assemblies were achieved through vigorous agitation (1,500 rpm) overnight at room temperature using a Benchmark Scientific MultiTherm shaker.

### Preparation of PSs via Flash Nanoprecipitation

PSs were formed using the confined impingement jets (CIJ) mixer described by Han et al. (*83*). BCP (either PPS-*b*-PEG-OMe, PPS-*b*-PEG-OH, or PPS-*b*-PEG-Phos) were dissolved in 500 μl of tetrahydrofuran (4.0% w/v) and aspirated via a 1 mL plastic disposable syringe. 500 μl of PBS was aspirated via a second 1 mL syringe. Utilizing the multiple impingement process described by Allen at al. (*28*), the two solutions were hand impinged against one another within the CIJ mixer. The supersaturated solution exited the mixer into an empty 20 mL glass scintillation vial. The solution, consisting of 20 mg of BCP and a 1:1 mixture of tetrahydrofuran and PBS, was evenly split between the two syringes and once again hand impinged against one another within the CIJ mixer. The process was repeated an additional three times, the final of which concluded by exiting the mixer into a 1.5 mL reservoir of PBS. Tetrahydrofuran was removed by placing the samples under vacuum for a minimum of six hours.

### Morphologic Confirmation via Cryogenic Transmission Electron Microscopy

Samples for cryogenic transmission electron microscopy (cryoTEM) were prepared by applying 3 µL of 10 mg/mL sample on pretreated lacey carbon 400 mesh TEM grids (Electron Microscopy Sciences). Following a 3 s blot, samples were plunge-frozen (Gatan Cryoplunge 3 freezer). Images of samples entrapped in vitreous ice were acquired using a field emission transmission electron microscope (JEOL 3200FS) operating at 300 keV with magnification ranging from 2,000 x to 12,000 x nominal magnification. Digital Micrograph software (Gatan) was used to align the individual frames of each micrograph to compensate for stage and beam-induced drift. Any further image processing conducted on the aligned frames was completed in ImageJ.

### Confirmation of Nanocarrier Morphology via Small Angle X-ray Scattering

Small angle X-ray scattering (SAXS) studies were performed at the DuPont-Northwestern-Dow Collaborative Access Team (DND-CAT) beamline at Argonne National Laboratory’s Advanced Photon Source (Argonne, IL, USA) with 10 keV (wavelength λ = 1.24 Å) collimated X-rays. All the samples were analyzed in the q-range (0.001 – 0.5 Å^− 1^), with a sample-to-detector distance of approximately 7.5 m and an exposure time of 3 s. The diffraction patterns of silver behenate were utilized to calibrate the q-range. The momentum transfer vector q is defined as q = 4π sinθ/λ, where θ is the scattering angle. Data reduction, consisting of the removal of solvent/buffer scattering from the acquired sample scattering, was completed using PRIMUS 2.8.2 software while model fitting was completed using SasView 4.0.1 software package. Nanocarrier morphology was confirmed by fitting the scattering profiles of PSs, FMs, and MCs with vesicle, flexible cylinder, and polymer micelle models, respectively.

### Spherical Nanocarrier Characterization via Dynamic Light Scattering (DLS)

The hydrodynamic diameter of each of the six spherical nanocarriers was obtained via DLS measurements recorded using a Zetasizer Nano (Malvern Instruments) equipped with a 4mW He-Ne 633 laser. The nanocarrier size distributions and number average diameters were acquired from measurements recorded from three independently formed formulations (n = 3). Samples were analyzed in 0.1x PBS prior to analysis.

### Zeta Potential Assessment via Electrophoretic Light Scattering (ELS)

The zeta potential of each of the nine nanocarriers was obtained using a Zetasizer Nano (Malvern Instruments). The zeta potential of each nanocarrier was averaged using measurements recorded from three independently formed formulations (n = 3). Samples were analyzed in 0.1× PBS prior to analysis.

### Nanocarrier Incubation with and Isolation from Human Plasma

Nanocarrier-protein isolation conditions were optimized as described in **Fig. S6-7**. Nanocarriers (5 mg/mL polymer concentration) of specified morphology and surface chemistry were incubated 1:1 with human plasma at 37 C, 220 rpm. This nanocarrier:protein ratio was determined to yield the formation of PCs having a protein concentration within the dynamic range of the Pierce 660nm assay (**Fig. S7**). Compared against common assays for the determination of protein concentration, the Pierce 660nm assay was determined to have the greatest specificity for protein with minimal interference by PEG-*b*-PPS polymer (**Fig. S6**). Nanocarrier-protein complexes were isolated by ultracentrifugation after incubation with human plasma for 2 h or 24 h. Ultracentrifugation was performed at 100,000 x g at 4 C for 45 min in an Optima MAX-XP ultracentrifuge (Beckman Coulter, Inc.). Supernatant was discarded, and nanocarrier-protein complexes were washed with PBS. This ultracentrifugation and washing process was repeated twice. After nanocarrier-protein complexes were isolated, the size and polydispersity were determined in three replicates by DLS and zeta potential was determined by ELS.

### Assessment of Total Protein Adsorption

Total adsorbed protein in a defined volume was determined by measuring protein concentration with the Pierce 660nm assay (ThermoFisher Scientific, Inc.) calibrated against a bovine serum albumin (BSA) concentration series. Measurements were made in three replicates. The concentration of adsorbed proteins was determined by **Eq. 1**:

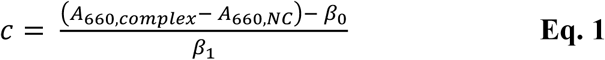

Where c is the determined protein concentration (in μg/mL), A_complex_ is the absorbance of 660 nm light (A_660_) measured in the NC-protein complex, A_NC_ is the A_660_ measured for the corresponding nanocarrier negative control lacking plasma, and β_0_ and β_1_ are the regression coefficients determined by simple linear regression.

### SDS-PAGE and Silver Staining

All protein samples were prepared in Laemmli buffer containing 10% 2-mercaptoethanol. 1 µg of each protein sample was separated in one dimension using 4-20% tris-glycine gels (Mini-PROTEAN TGX, Bio-Rad Laboratories, Inc.). A 1:6 dilution of the PageRuler Plus pre-stained protein ladder (10 µl) was used as a molecular weight standard (ThermoFisher Scientific, Inc.). BSA and human plasma samples were included on each gel for comparison. Electrophoresis proceeded at 110 V for 70 minutes. Gels were washed twice with MQ water, then subsequently fixed overnight in a 30% ethanol, 10% acetic acid solution (6:3:1 water:ethanol:acetic acid). On the following day, gels were washed twice in 10% ethanol, followed by two washing steps in water. The gels were subsequently silver stained in clean glass trays using the Pierce Silver Stain kit for mass spectrometry (ThermoFisher Scientific, Inc.). Each gel was developed for approximately 45 seconds prior to halting stain development in 5% acetic acid. The gels were briefly washed in MQ water prior to imaging. Grayscale TIFF images (800 dpi) were obtained for each gel using an inexpensive Epson V39 scanner (Seiko Epson, Corp.). Gel images were cropped and straightened in ImageJ (*84*).

### Protein identification and label-free quantification

Prior to protein identification, protein samples were cleaned up by SDS-PAGE following established procedures. A total of 5 μg of adsorbed protein was prepared in Laemelli buffer and β-mercaptoethanol, loaded into a 4-20% Tris-glycine polyacrylamide gel, and were briefly electrophoresed at 120 V for roughly 5 minutes to achieve 3-5 mm separation from the well. A 1 cm x 1 cm band containing the mixed adsorbed protein population was excised and processed further for label free quantification. After in-gel trypsin digestion, samples were analyzed using a Q Exactive HF Hybrid Quadrupole-Orbitrap mass spectrometer (ThermoFisher Scientific). Data was collected from two technical replicates. The data was searched against the UniProt *Homo sapiens* database using the MaxQuant search engine and label-free quantification (LFQ) was performed based on the MS1 peptide intensity. Proteins were identified using a threshold of >1 unique peptide and a 1% FDR. A total of 171 proteins were identified and quantified by these procedures.

### Statistical analysis of proteomic data

The LFQ intensity was log2 transformed and normalized by median subtraction. Imputation was applied to fill in missing values based on normal distribution, which is a recommended procedure for LFQ analysis. Statistically significant differences in protein relative abundance across the nine nanocarrier formulations were determined by ANOVA. Within each formulation, significant differences in protein relative abundance at 2 h versus 24 h were determined by pairwise t-tests. The p-values were calculated by FDR-based permutation procedure.

### Network and biological process enrichment analysis

Gene ontology (GO) analyses were performed on the subset of 121 quantified proteins, determined to be significantly enriched across the nine nanocarrier formulations at 2h. A simplified analysis of biological process GO slim terms was initially presented. A more comprehensive of enriched biological process GO terms was performed using BiNGO (*85*). Overrepresented biological process terms were determined against the *H. sapiens* reference set using the hypergeometric test corrected using the Benjamini and Hochberg False Discovery Rate (FDR) with a 1e-20 significance level. A highly stringent p-value was used to focus on the most significant subset of enriched terms and to produce a network that is interpretable. Network diagrams were constructed in Cytoscape 3.8.0.

### Endotoxin Testing

Nanocarrier formulations were tested for endotoxin contamination using the QUANTI-Blue (InvivoGen) colorimetric enzyme assay. Raw-Blue cells (InvivoGen) were plated in 48 well plates (2.5 × 10^5^ cells/mL, 400 μl). Subsequently, 40 μl of each nanocarrier formulation (5 mg/mL) was added per well and incubated for 24 h at 37°C with a 5% CO_2_ atmosphere. Upon completion, 20 μl of cell supernatant was collected and added to 200 μl of the QUANTI-Blue detection media. Cell supernatants were incubated with the detection media for an additional 18 h at 37°C. SEAP activity was assessed by taking the optical density (OD) at 625 nm with a microplate reader (SpectraMax M3, Molecular Devices).

### Assessment of Nanocarrier-Induced Cytokine Secretion

Whole blood samples containing ethylenediaminetetraacetic acid (EDTA) collected from three individual, healthy donors were acquired from Research Blood Components, LLC. Nanocarrier formulations (10 mg/mL, 8 μl) were added to 152 μl of human whole blood from each of the donors for 4 or 20 h at 37°C with a 5% CO_2_ atmosphere. Lipopolysaccharide (LPS) (InvivoGen) and R848 (InvivoGen) were included in the analysis as positive controls and were tested at final concentrations of 1.0 and 10 μg/mL, respectively. After incubation, plasma was collected via centrifugation. Samples were subsequently prepared and analyzed following the manufacturer’s protocol. Data acquisition was completed using a BD LSRFortessa. Acquired data was analyzed using the online Cytobank analysis suite.

### Assessment of Nanocarrier-Induced Complement Activation

Nanocarrier formulations (10 mg/mL, 5 μl) were added to 95 μl of pooled complement human serum (Innovative Research) and incubated for 1 h at 37°C while being mixed at 200 rpm. Upon completion, samples were treated with 5 mM EDTA to stop the activation of complement and were placed on ice until further use. The concentrations of the human anaphylatoxins C3a, C4a, and C5a were quantitatively assessed using a BD™ CBA Human Anaphylatoxin Kit. Generation of the C3a, C4a, and C5a standards and sample preparation were conducted following the manufacturer’s protocol. Data acquisition was completed using a BD LSRFortessa. Acquired data was analyzed using the online Cytobank analysis suite.

### THP-1 Culture Conditions

THP-1 cells (ATCC), a human leukemic monocyte cell line, were cultured in Roswell Park Memorial Institute (RPMI)-1640 medium (Gibco) supplemented with 10% fetal bovine serum (FBS) (Sigma-Aldrich), 1% GlutaMax (Gibco), and 1% penicillin– streptomycin (Gibco), hereafter referred to as complete RPMI. Culture conditions were maintained at 37°C with a 5% CO_2_ atmosphere. Cells were subcultured as needed to insure cell concentration remained below 1.0 x 10^6^ cells/mL. Cells used for cytotoxicity or nanocarrier uptake studies were collected between passages 5 and 10.

### THP-1 Differentiation to Macrophage-like Phenotype

THP-1 monocytes were differentiated toward a macrophage-like phenotype (*86*). THP-1 cells, suspended in complete RPMI at a concentration of 2.0 x 10^5^ cells/mL, were treated with phorbol 12-myristate 13-acetate (PMA) at a final concentration of 25 nM upon dilution. Cells were plated in 48 well plates (2.0 x 10^5^ cells/mL, 500 μL) and incubated at 37°C with a 5% CO_2_ atmosphere for 48 h. Following differentiation, PMA-containing media was removed and replaced with complete RPMI. THP-1 differentiated macrophages were incubated for an additional 24 h in PMA-free media prior to being washed and used for subsequent experiments.

### Immature Dendritic Cell Differentiation from Human Monocytes

Monocyte derived immature dendritic cells were obtained by following a previously published protocol (*87*). Negatively selected human monocytes (Astarte Biologics) were suspended in RPMI 1640 supplemented with 2% human AB serum (Sigma-Aldrich), 1% GlutaMax (Gibco), 1% penicillin– streptomycin (Gibco), 1000 U/mL of Granulocyte-macrophage colony-stimulating factor (GM-CSF) (Shenandoah Biotechnology), and 500 U/mL of IL-4 (Shenandoah Biotechnology), hereafter referred to as complete DC media. Monocytes were plated in 6 well plates (5.0 x 10^5^ cells/mL, 3.0 mL) and maintained at 37°C with a 5% CO_2_ atmosphere for 5 days. Three days after plating, half of the media volume was collected, centrifuged at 500xg for 5 min, and discarded. The cell pellet was resuspended in an equivalent volume of fresh complete DC media and redistributed among the wells.

### Assessment of Nanocarrier Cytotoxicity

The 3-(4,5-Dimethylthiazol-2-yl)-2,5-Diphenyltetrazolium Bromide) (MTT) assay was used to quantify the percentage of metabolically active cells as a readout of cell viability. THP-1 monocytes (5.0 × 10^5^ cells/mL, 200 μl) were plated in U-bottom 96-well plates. Nanocarriers (20 μl) at concentrations of 0, 1, 2, 5 and 10 mg/mL (prepared in PBS) were added to the wells in quadruplicate and incubated for 24 h. 20 μl of MTT (5 mg/mL prepared in PBS) was added to each well and incubated for an additional 6 h. Plates were centrifuged (500 x g, 5 min) before removing supernatant from each well. Subsequently, dimethyl sulfoxide (200 μl) was added to each well to dissolve deposited formazan crystals. Absorbance measurements at 570 nm were acquired using a microplate reader (SpectraMax M3, Molecular Devices). Cell viability was assessed through **Eq. 2**:

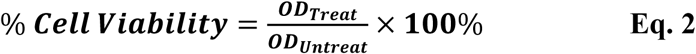

Where OD_Treat_ is the optical density of a nanocarrier treated sample and OD_Untreat_ is the optical density of an untreated sample. The average percentage of viable cells was calculated from eight total measurements (n = 8) acquired through two independent experiments.

### Cellular Uptake of Nanocarriers *in vitro*

All nanocarrier uptake studies were conducted utilizing a 4 h incubation period in serum-free media. Nanocarriers were tested under three conditions: (i) pristine (no exposure to pooled human plasma; BCP concentration of 5 mg/mL), (ii) following a 2 h incubation with pooled human plasma (1:1 volumetric ratio of nanocarrier formulation to human plasma; BCP concentration of 5 mg/mL), or (iii) following a 24 h incubation with human plasma (1:1 volumetric ratio of nanocarrier formulation to human plasma; BCP concentration of 5 mg/mL). Pooled human plasma was acquired from Zen-Bio Inc. and contained acid citrate dextrose (ACD) as an anti-coagulant.

THP-1 monocytes were collected via centrifugation (500 x g, 5 min), washed with serum-free RPMI 1640, and recentrifuged. Pelleted cells were resuspended in serum-free RPMI 1640 to a final concentration of 2.5 × 10^5^ cells/mL and plated in 48 well plates (400 μl). Cells were incubated for 1 h at 37°C with a 5% CO_2_ atmosphere prior to the introduction of nanocarriers. After 1 h, 40 μl (5 mg/mL) of each nanocarrier formulation for each defined condition were added to individual wells. Cells were incubated for 4 h at 37°C with a 5% CO_2_ atmosphere following the addition of the nanocarrier formulations before being collected for uptake analysis via flow cytometry. Cells were stained with Zombie Aqua cell viability dye (BioLegend) for 15 min at a 1:200 dilution in cell staining buffer (BioLegend), washed with cell staining buffer, and briefly fixed with 2% paraformaldehyde prior to analysis on a BD LSRFortessa.

For THP-1 differentiated macrophages, complete RPMI was removed and the adherent cells were washed with serum-free RPMI 1640. Fresh serum-free RPMI (400 μl) was added to each well. Cells were incubated for 1 h at 37°C with a 5% CO_2_ atmosphere prior to the introduction of nanocarriers. After 1 h, 40 μl (5 mg/mL) of each nanocarrier formulation for each defined condition were added to individual wells. Cells were incubated for 4 h at 37°C with a 5% CO_2_ atmosphere following the addition of the nanocarrier formulations before being collected for uptake analysis via flow cytometry. Cells were stained with Zombie Aqua cell dye (BioLegend) for 15 min at a 1:200 dilution in cell staining buffer (BioLegend), washed with cell staining buffer, and briefly fixed with 2% paraformaldehyde prior to analysis on a BD LSRFortessa.

After 5 days of culture in complete DC media, differentiated immature dendritic cells were collected via centrifugation (500 x g, 5 min), washed with serum-free RPMI 1640, and recentrifuged. Pelleted cells were resuspended in serum-free RPMI 1640 to a final concentration of 2.5 × 10^5^ cells/mL and plated in 48 well plates (300 μl). Cells were incubated for 1 h at 37°C with a 5% CO_2_ atmosphere prior to the introduction of nanocarriers. After 1 h, 30 μl (5 mg/mL) of each nanocarrier formulation for each defined condition were added to individual wells. Cells were incubated for 4 h at 37°C with a 5% CO_2_ atmosphere following the addition of the nanocarrier formulations before being collected for uptake analysis via flow cytometry. Cells were stained with Zombie Aqua cell viability dye (BioLegend) for 15 min at a 1:200 dilution in cell staining buffer (BioLegend) and subsequently stained with an antibody mixture containing allophycocyanin (APC) anti-human CD209 (DC-SIGN) and fluorescein isothiocyanate (FITC) anti-human CD16 for 20 min. Cells were subsequently washed with cell staining buffer and fixed with 2% paraformaldehyde prior to analysis on a BD LSRFortessa. Acquired flow cytometry data for all cell types was analyzed using the online Cytobank analysis suite.

## Acknowledgements

The authors acknowledge Jonathan Remis (Structural Biology Facility, NU) for assisting with cryoTEM and Dr. Matt Clutter (High Throughput Analysis Laboratory, NU) for assisting with the acquisition of data used in the nanocarrier immunogenicity and complement activation analyses. The authors acknowledge Dr. Young Ah Goo (NU Proteomic Facility) and Dr. Byoungkyu Cho (NU Proteomics Facility) for their support and assistance with the proteomic data acquisition.

## Author contributions

N.B.K., M.P.V., and E.A.S. contributed to the conception and study design. N.B.K. and Y.Y. synthesized and characterized materials. N.B.K. and M.P.V. contributed to the preparation and physicochemical characterization of the nanocarriers. N.B.K., S.B., and M.P.V. contributed to SAXS characterization of nanocarriers and corresponding analysis. N.B.K., M.P.V., S.D.A., S.Y. contributed to experiments performed *in vitro*. M.P.V. and E.A.S. contributed to the conception and experimental design for nanocarrier protein corona isolation, protein quantification, protein identification, and analysis. M.P.V. and N.B.K. performed the protein adsorption experiments. M.P.V. performed the proteomic experiments and the corresponding analyses. M.P.V., N.B.K., S.B., and M.A.F., contributed to the supplementary experiments. N.B.K., M.P.V., S.D.A., and E.A.S. contributed to the statistical analysis. M.P.V. and S.D.A. contributed to the graphical design of the illustrations. M.P.V, N.B.K., and E.A.S. contributed to writing the manuscript and to the preparation of figures.

## Data and materials availability

Materials and data will be made available upon request.

