## Supplemental data for "The Combination of Morphology and Surface Chemistry Defines the Biological Identity of Nanocarriers in Human Blood"

**Affiliations:**
[1]Department of Biomedical Engineering, Northwestern University, Evanston, Illinois, USA
[2]Interdisciplinary Biological Sciences, Northwestern University, Evanston, Illinois, USA
[3]Chemistry of Life Processes Institute, Northwestern University, Evanston, Illinois, USA
[4]Simpson Querrey Institute, Northwestern University, Chicago, Illinois, USA
[5]Robert H. Lurie Comprehensive Cancer Center, Northwestern University, Chicago, Illinois, USA

[†]These authors are co-first authors and contributed equally to this work

*To whom correspondence should be addressed:
Evan Scott, PhD
Assistant Professor of Biomedical Engineering
Northwestern University
2145 Sheridan Road
Evanston, IL 60208


### Supplementary Figures

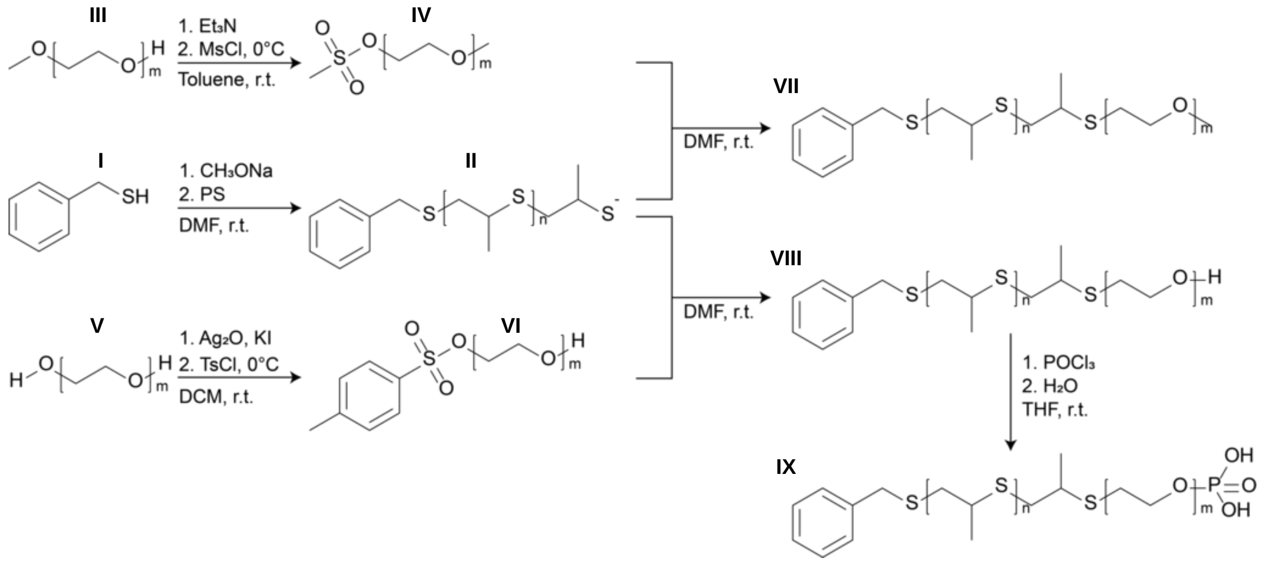


**Figure S1. Synthetic pathway used to generate methoxy, hydroxyl, or phosphate terminated BCPs.** Benzyl mercaptan (**I**) is base-activated and employed to perform anionic ring-opening polymerization on propylene sulfide to produce poly(propylene sulfide) homopolymer (**II**). To function as end capping agents for **II**, mPEG (**III**) or PEG (**V**) are used to prepare mPEG-mesylate (**IV**) or HO-PEG-tosylate (**VI**). These two sulfonate PEG derivates are used to end cap **II**, resulting in Bn-PPS-*b*-PEG-OMe (**VII)** and Bn-PPS-*b*-PEG-OH (**VIII**), respectively. The terminal hydroxyl on **VIII** is subsequently converted into a phosphate group to form Bn-PPS-*b*-PEG-Phos (**IX**).

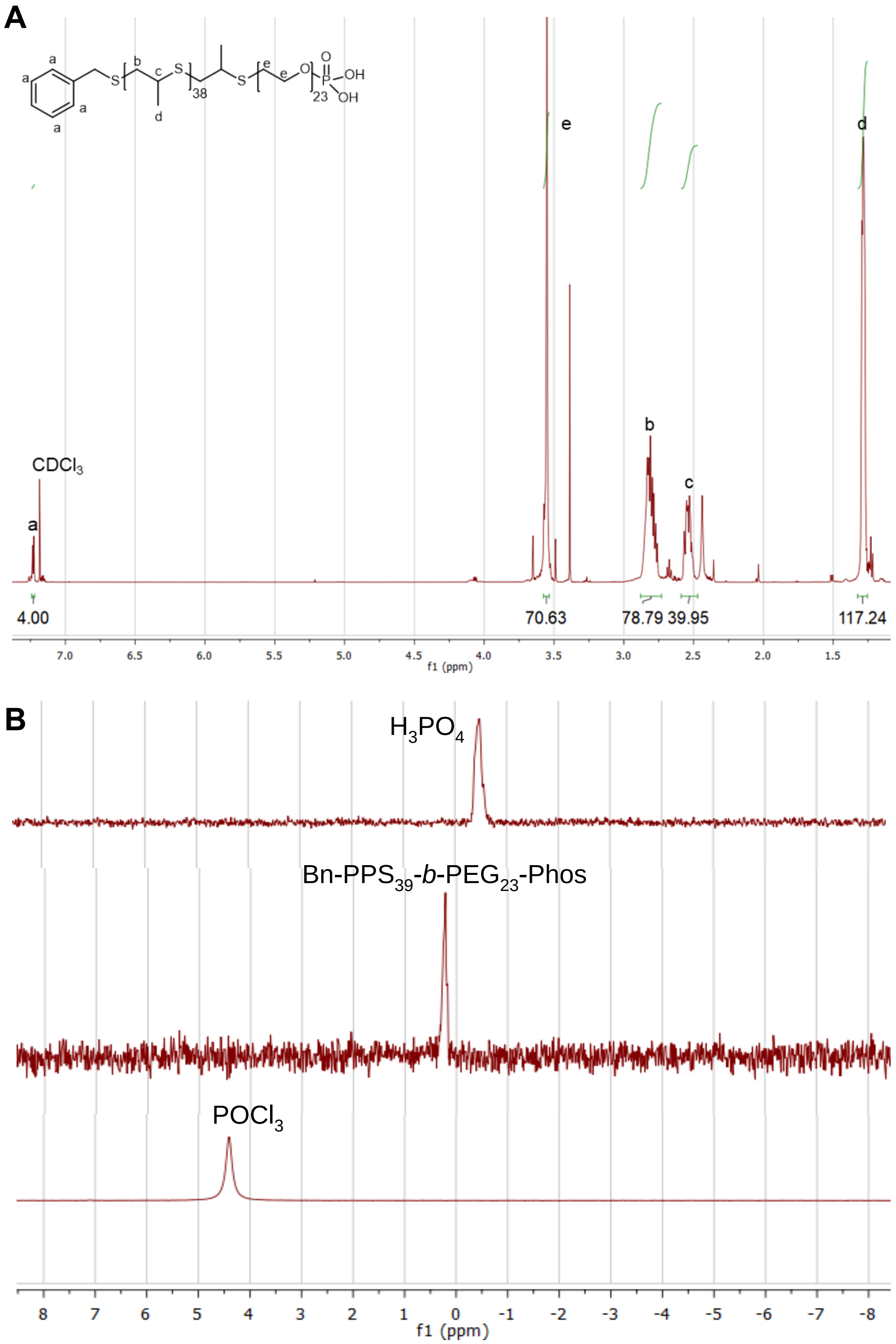


**Figure S2. Representative NMR of Bn-PPS-*b*-PEG-Phos.** (**A**) Representative $^{1}$H NMR spectrum for Bn-PPS$_{39}$-*b*-PEG$_{23}$-Phos. (**B**) Representative $^{31}$P NMR spectrum for Bn-PPS$_{39}$-*b*-PEG$_{23}$-Phos. The rightward shift of the phosphorous peak toward 0 ppm signifies the successful functionalization of the BCP.

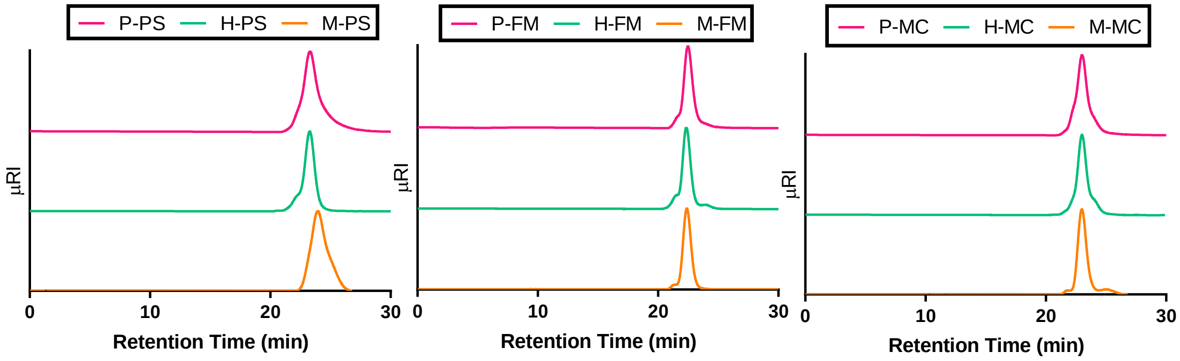


**Figure S3. Gel permeation chromatography (GPC) chromatograms of synthesized BCPs.** Overlay of chromatograms for Phos (P)-, OH (H)-, and MeO (M)-functionalized BCPs that assemble into PSs (left), FMs (middle), and MCs (right).

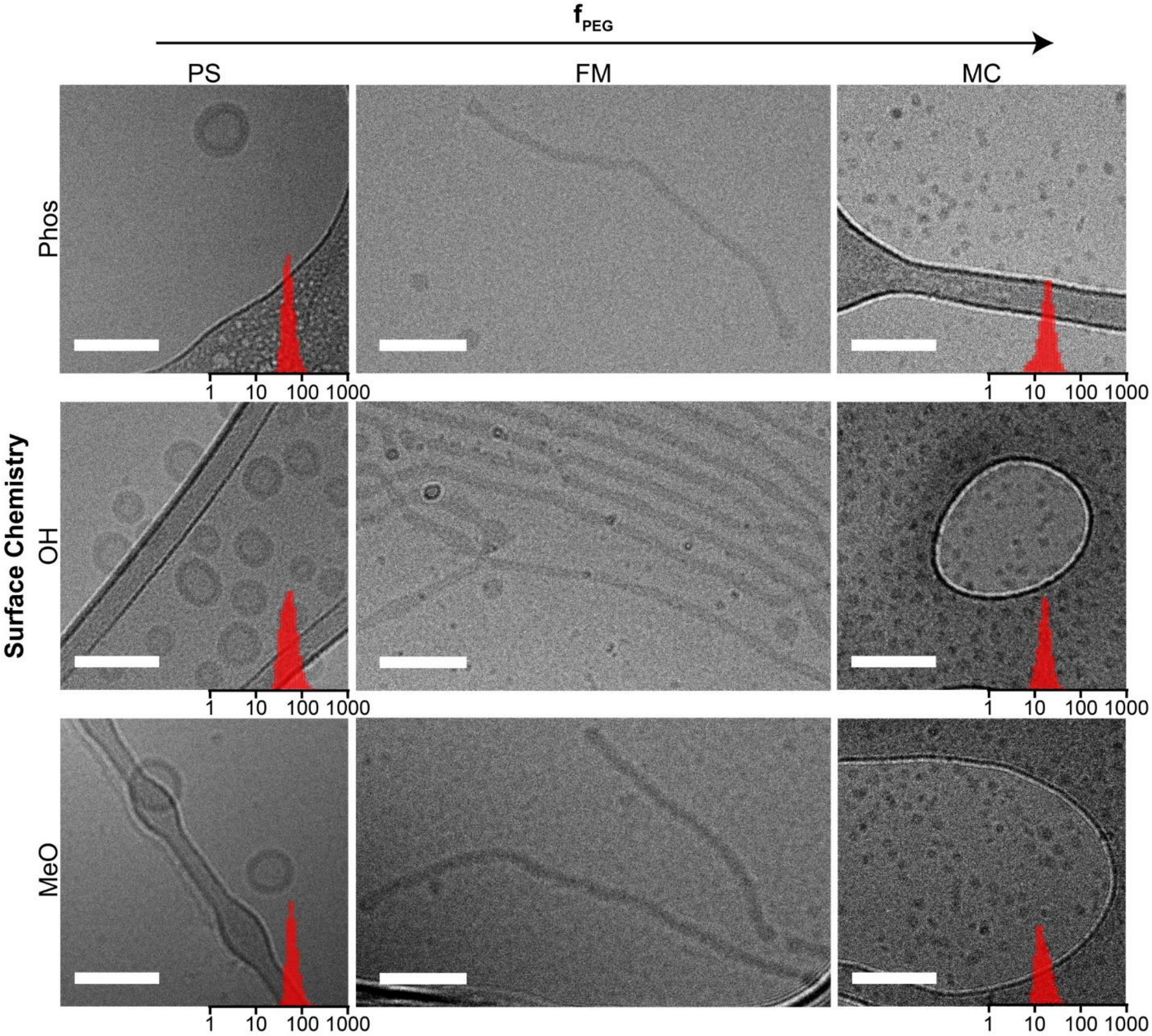


**Figure S4. Morphological characterization of PEG-*b*-PPS nanocarriers by CryoTEM.** For the spherical nanocarriers, PS and MC, the CryoTEM micrographs are displayed with overlaid histograms of particle hydrodynamic diameters (determined by DLS). FM micrographs are re-displayed here from the main text for the purpose of comparison. Scale bar = 100 nm.

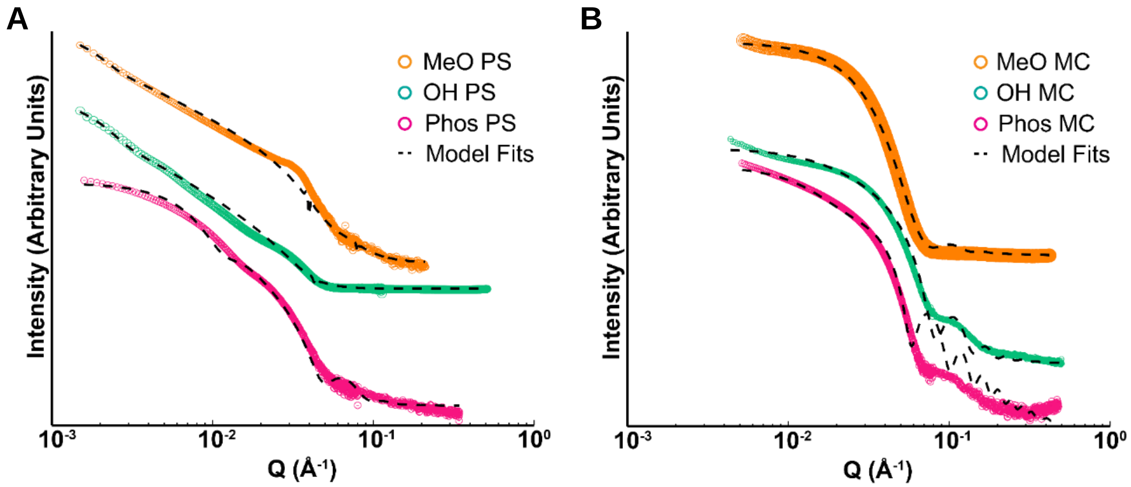


**Figure S5. SAXS profiles of PEG-*b*-PPS polymersomes and micelles overlaid with corresponding model fits.** (**A**) SAXS profiles of PEG-b-PPS PSs overlaid with corresponding fit for a vesicle model ($\chi 2$ = 0.555, 0.363, 0.0645 for MeO-, OH-, and Phos-PSs, respectively). (**B**) SAXS profiles of PEG-b-PPS MCs overlaid with corresponding fit for a polymer micelle model ($\chi 2$ = 0.0047, 0.0017, 0.0084 for MeO-, OH-, and Phos-MCs, respectively).

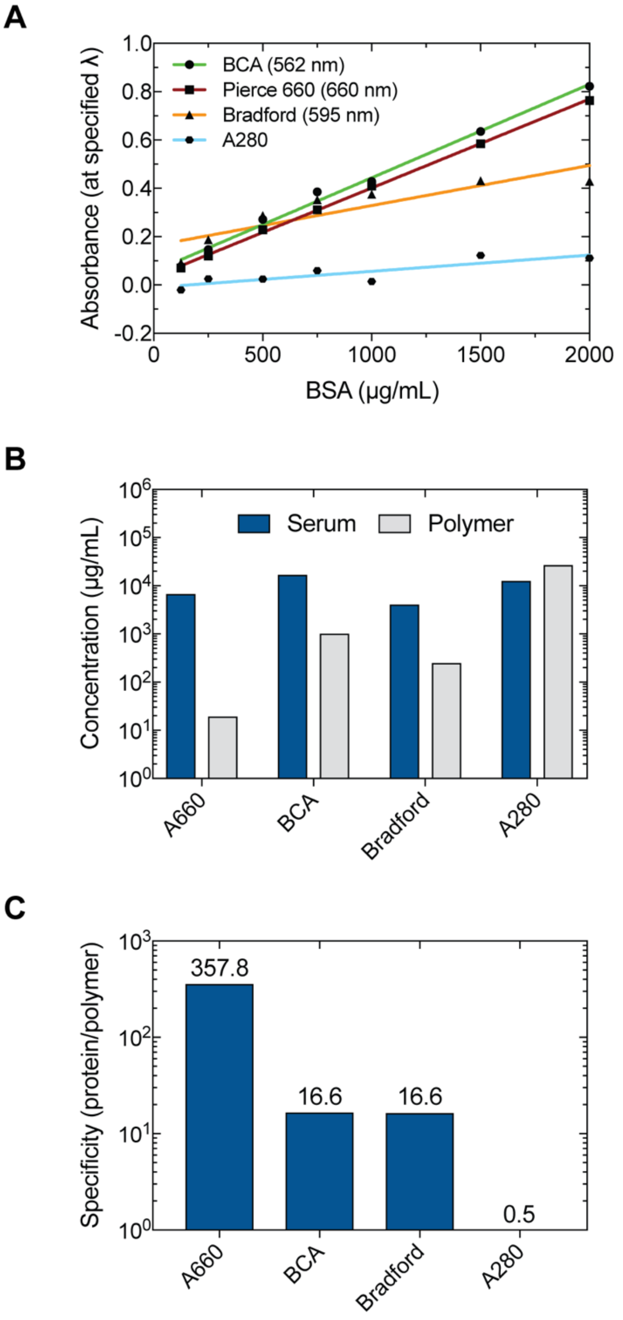


**Figure S6. The Pierce A660 assay determines protein concentrations with minimal interference from PEG-*b*-PPS polymer.** (**A**) BSA calibration curves for each assay. A660: Pierce 660 nm reagent ($y = 3.7e\text{-}4x + 3.4e\text{-}2$; $r^2 = 0.99$; F=5292, p<0.0001); BCA: Bicinchoninic acid assay (562 nm; $y = 3.9e\text{-}4x + 5.6e\text{-}2$; $r^2 = 0.76$; F=666.7, p<0.0001); Bradford assay (595 nm; $y = 1.7e\text{-}4x + 1.6e\text{-}1$; $r^2 = 0.81$; F=17.8, $p<0.01$); A280: concentration determination by absorbance of 280 nm light ($y = 6.71e\text{-}5x - 1.0e\text{-}2$; $r^2 = 0.76$; F=15.88, $p<0.05$). (**B**) Serum and Polymer are Fetal Bovine Serum (FBS; 14.5 mg/mL) and PEG-*b*-PPS methoxy-polymersome polymer (5 mg/mL), respectively. (**C**) Assay specificity for protein, calculated as the ratio of protein/polymer for the protein concentrations determined in (**B**).

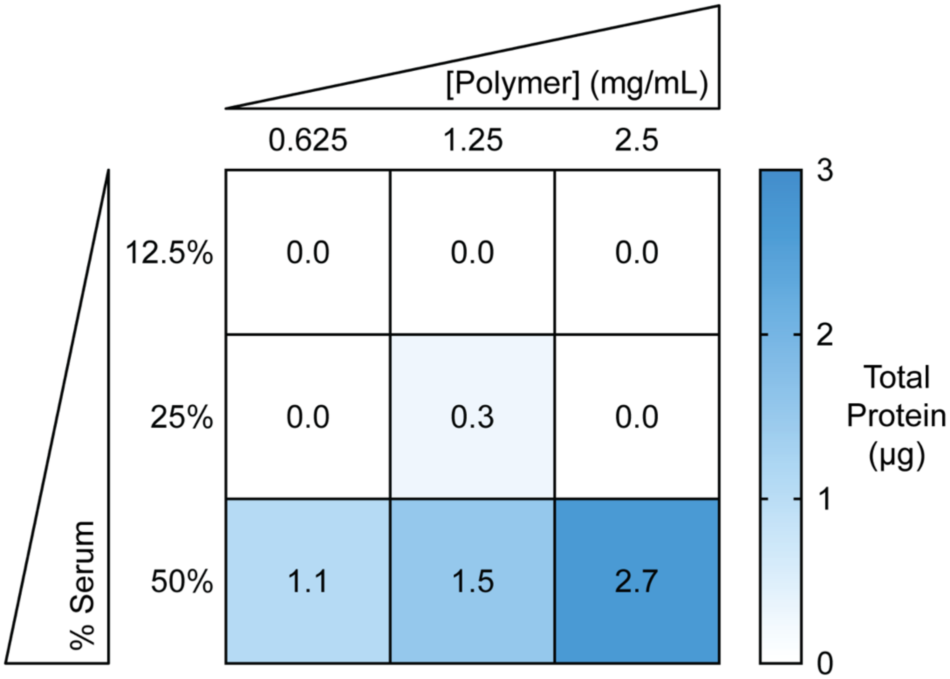


**Figure S7. Determination of the protein to polymer ratio for use in the protein adsorption studies.** For method development purposes, MeO PS were incubated with fetal bovine serum for 2 h under the specified serum and NC polymer concentrations. NC-protein complexes were isolated after multiple rounds of centrifugation at 100,000 x g for 45 min at 4 °C and PBS washing. The average total protein recovered in a 200 µl volume is shown. Protein concentration was determined using the A660 assay.

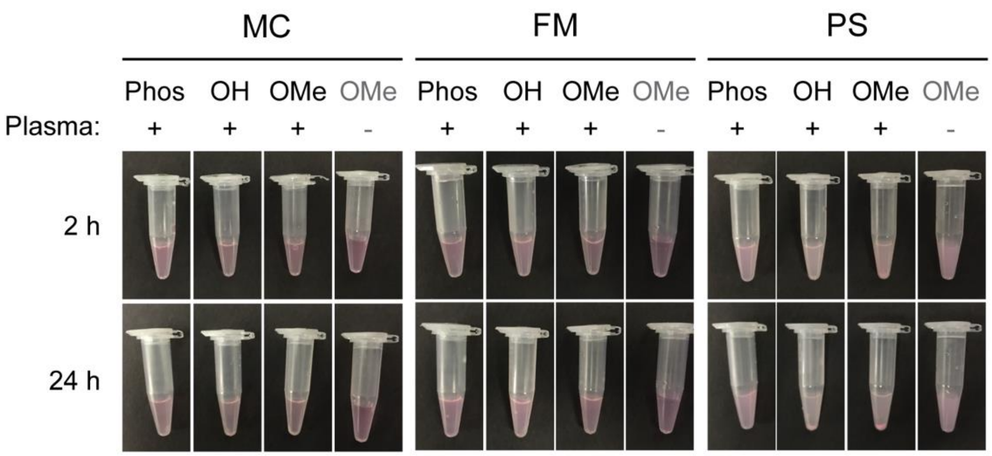


**Figure S8. Assessment of NC agglomeration in the presence of human plasma proteins.** Images of NCs incubated for 2 h or 24 h in the presence or absence of plasma.

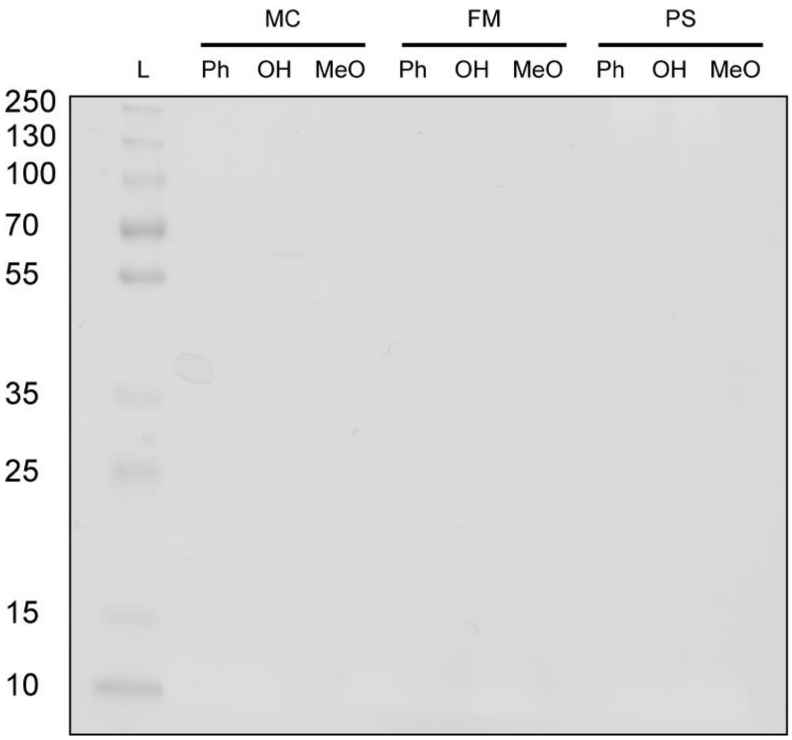


**Figure S9. SDS-PAGE of nanocarrier negative controls used in the protein adsorption studies.** The specified PEG-*b*-PPS NC (5 mg/mL polymer concentration) was incubated 1:1 in 1x PBS and incubated at 37 °C, 220 rpm for 24 h. NCs were isolated by ultracentrifugation, prepared in Laemmli buffer containing 2-mercaptoethanol, loaded into a 12% tris-glycine SDS-PAGE gel.

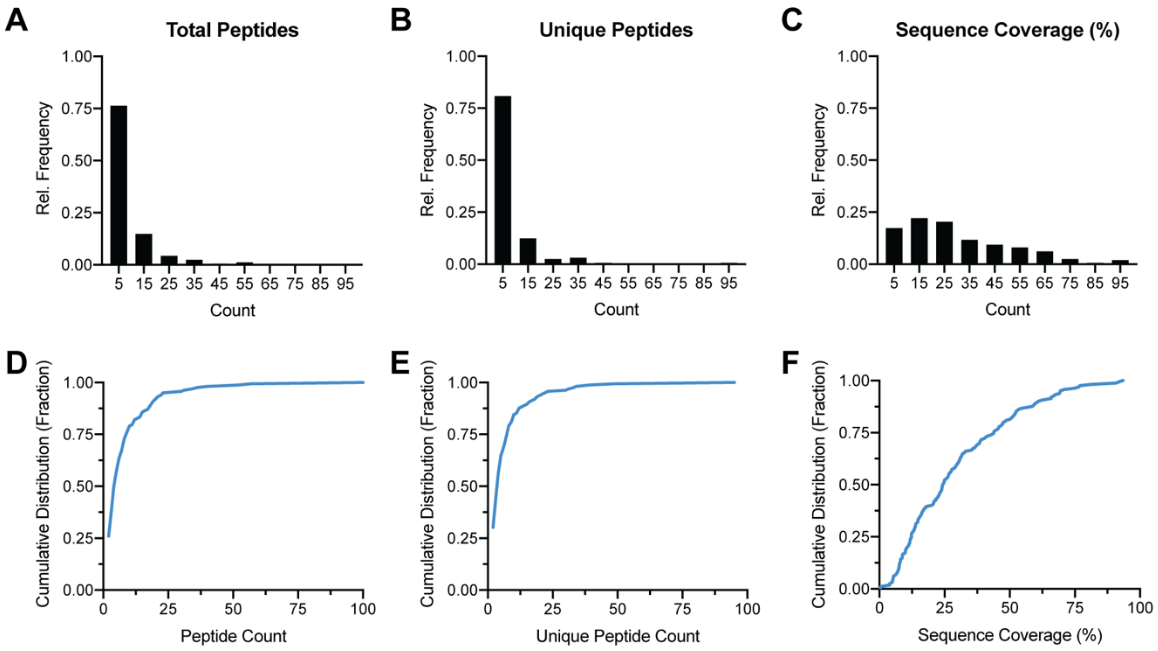


**Figure S10. Summary of protein identification statistics.** (**A-C**) Histograms showing the relative frequency of the (**A**) total peptides, (**B**) unique peptides, and (**C**) sequence coverage (%) for peptides used to identify proteins in the proteomic analyses (bin width = 10; bin center values are shown on the x-axis). (**D-F**) Cumulative frequency distributions for the (**D**) total peptides, (**E**) unique peptides, and (**F**) sequence coverage (%).

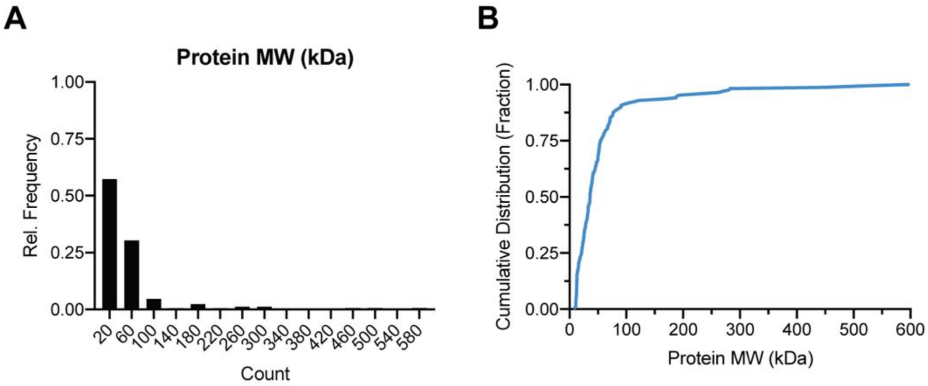


**Figure S11. Molecular weight distribution of identified human plasma proteins in this study.** (**A**) Histogram displaying the molecular weight (kDa) distribution of the 171 identified plasma proteins (bin width = 40; bin center values are shown on the x-axis). (**B**) Cumulative distribution of protein molecular weight.

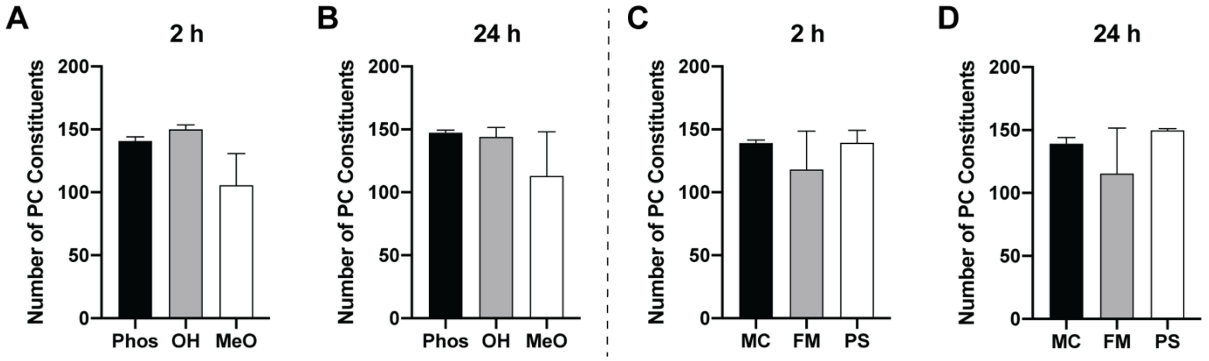


**Figure S12. The number of protein corona (PC) constituents in bulk surface chemistry and morphological groupings.** (**A, B**) The average number of PC constituents in nanocarriers pooled by surface chemistry at 2 h (**A**) and 24 h (**B**). (**C, D**) The average number of PC constituents in nanocarriers pooled by morphology at 2 h (**C**) and 24 h (**D**). Groupings were defined using the mean number of protein constituents determined in individual NC formulations. All pairwise comparisons were determined to be not significant, as determined by ANOVA with Tukey's multiple comparisons test (5% significance level). In all cases, error bars represent s.e.m.

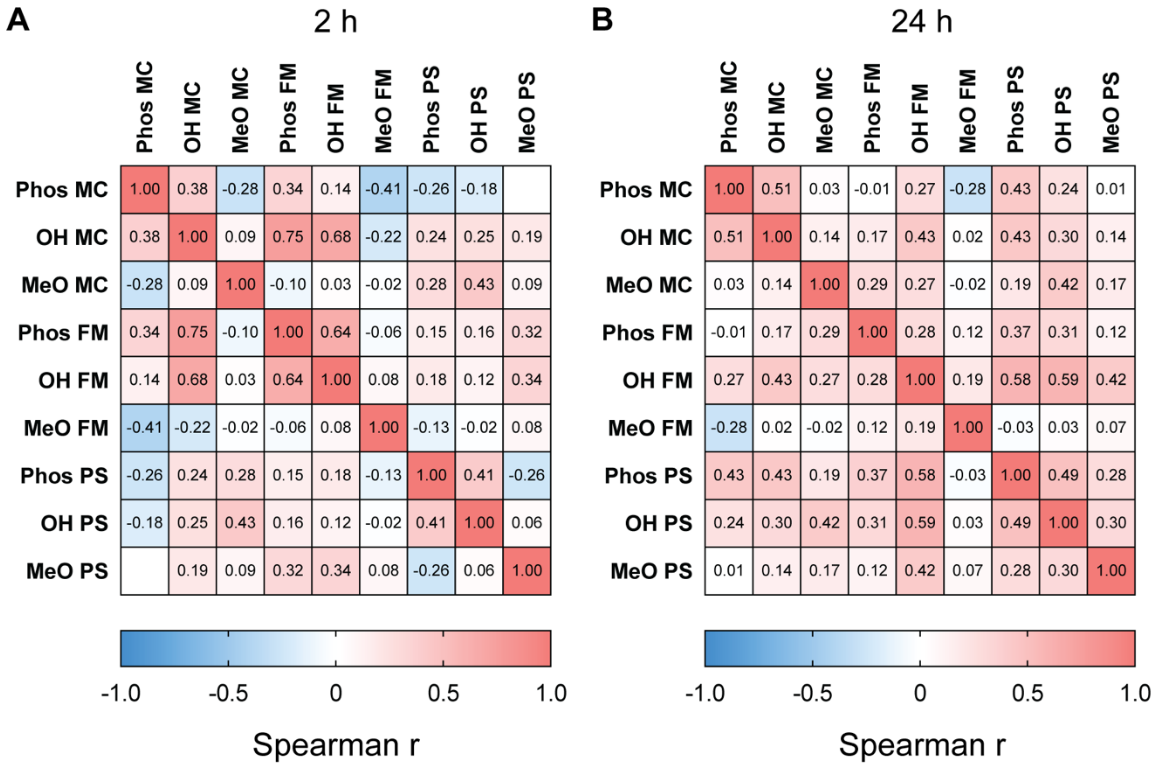


**Figure S13. Spearman correlation matrix for the complete set of nanocarrier protein corona constituent relative abundances.** The spearman correlation coefficient (r) was calculated for the complete set of 171 identified proteins at (**A**) 2 h and (**B**) 24 h. Blank cells have r<0.01.

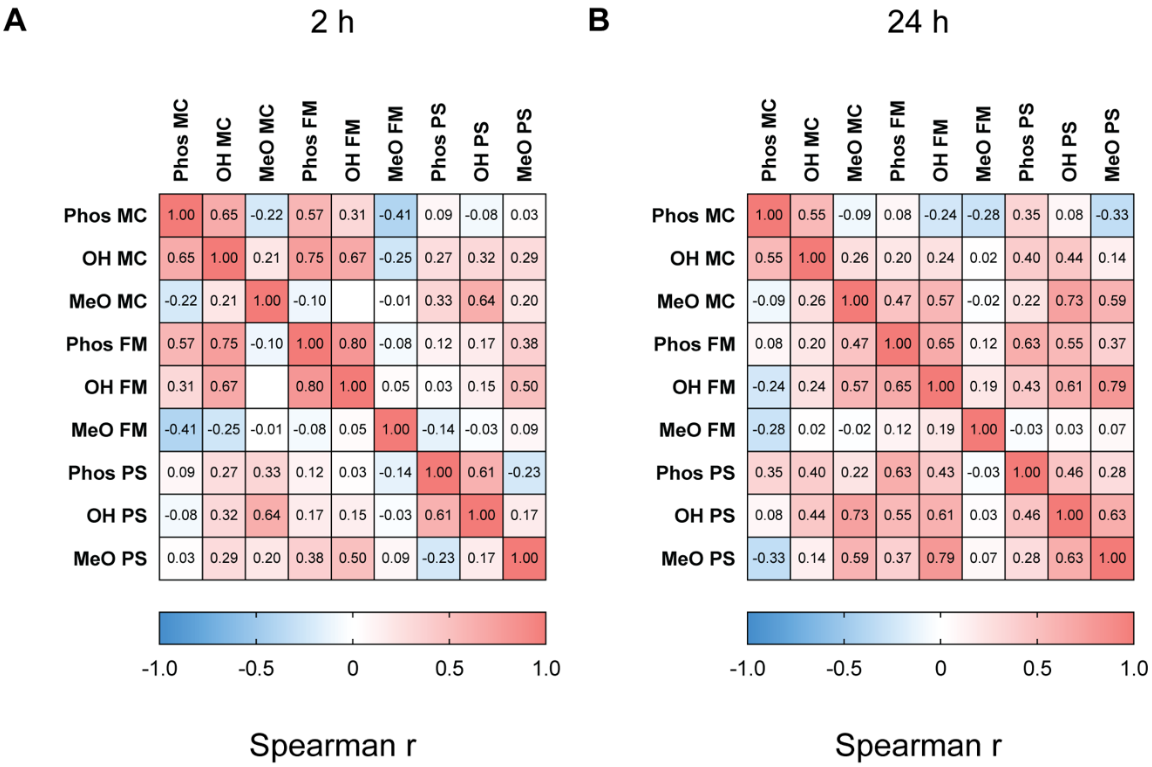


**Figure S14. Spearman correlation matrix for the shared set of nanocarrier protein corona constituent relative abundances.** The spearman correlation coefficient (r) was calculated for the shared set of identified proteins at (**A**) 2 h and (**B**) 24 h. Proteins that were not detected in all nine NC protein coronas were excluded from this particular analysis. Blank cells have r<0.01.

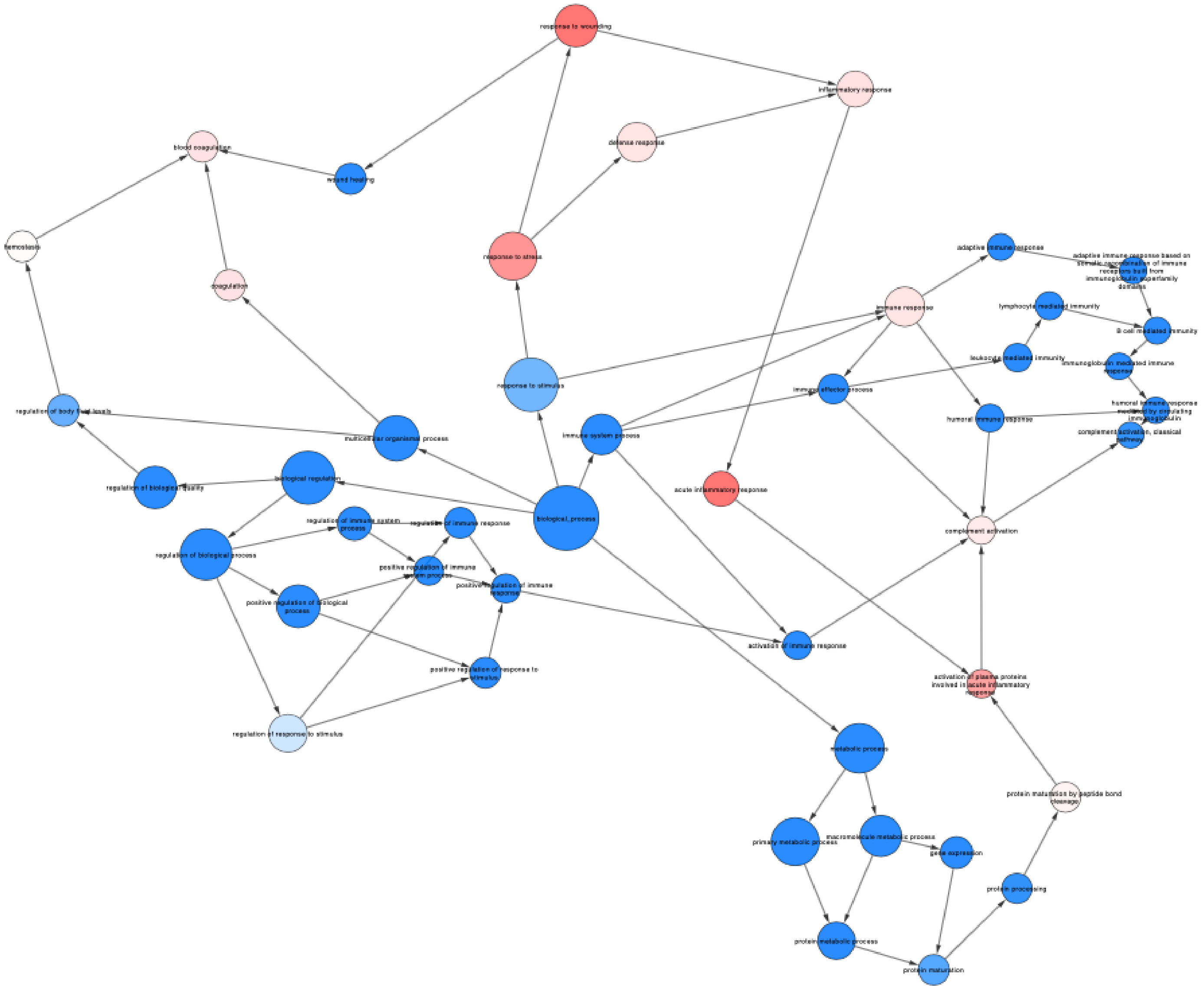


**Figure S15. Full annotations for the 2 h biological process gene ontology network.** For the purpose of completeness, the full annotations are displayed for the 2 h network of biological process gene ontology terms presented in the main text.

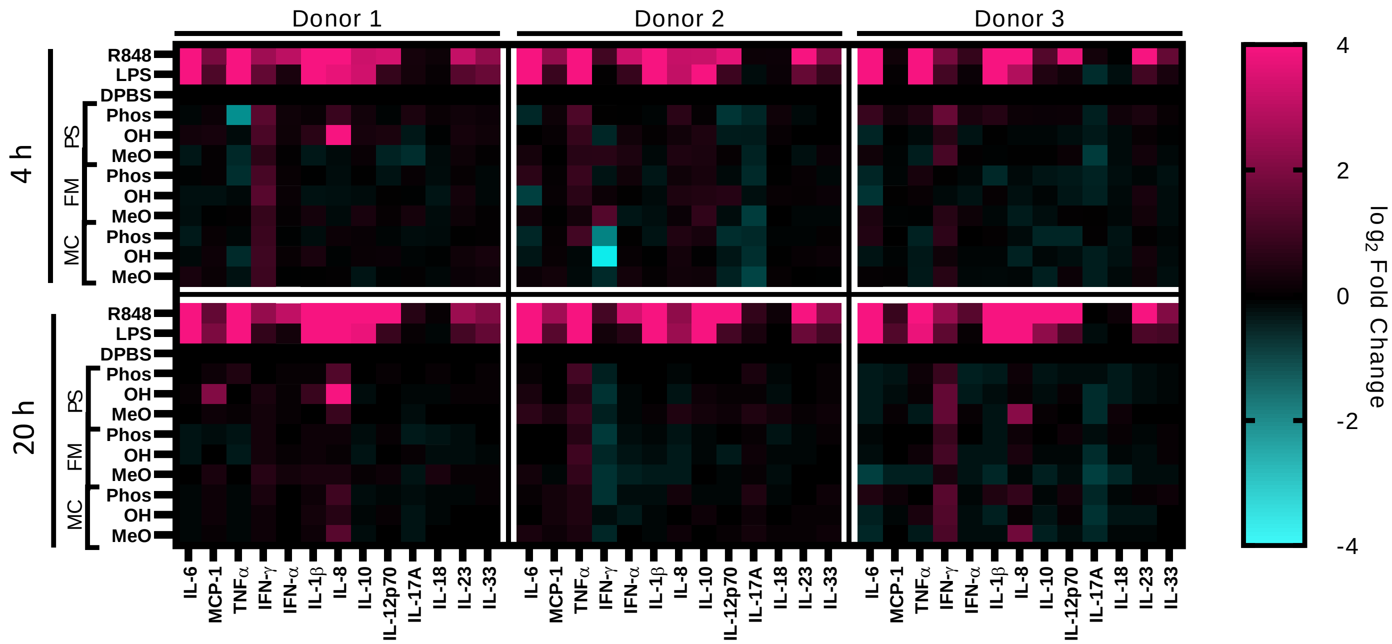


**Figure S16. Individual donor cytokine profiles following 4 h and 20 h incubations of nanocarrier formulations with whole blood.** The upper row depicts the donor cytokine profiles following a 4 h incubation of donor whole blood with the various nanocarrier formulations. The lower row depicts the donor cytokine profiles following a 20 h incubation of donor whole blood with the various nanocarrier formulations. Individual donors (n = 1) are split into columns: Donor 1 (left), Donor 2 (middle), and Donor 3 (right). The 4 h data is re-displayed from Figure 4 for the purpose of comparison.

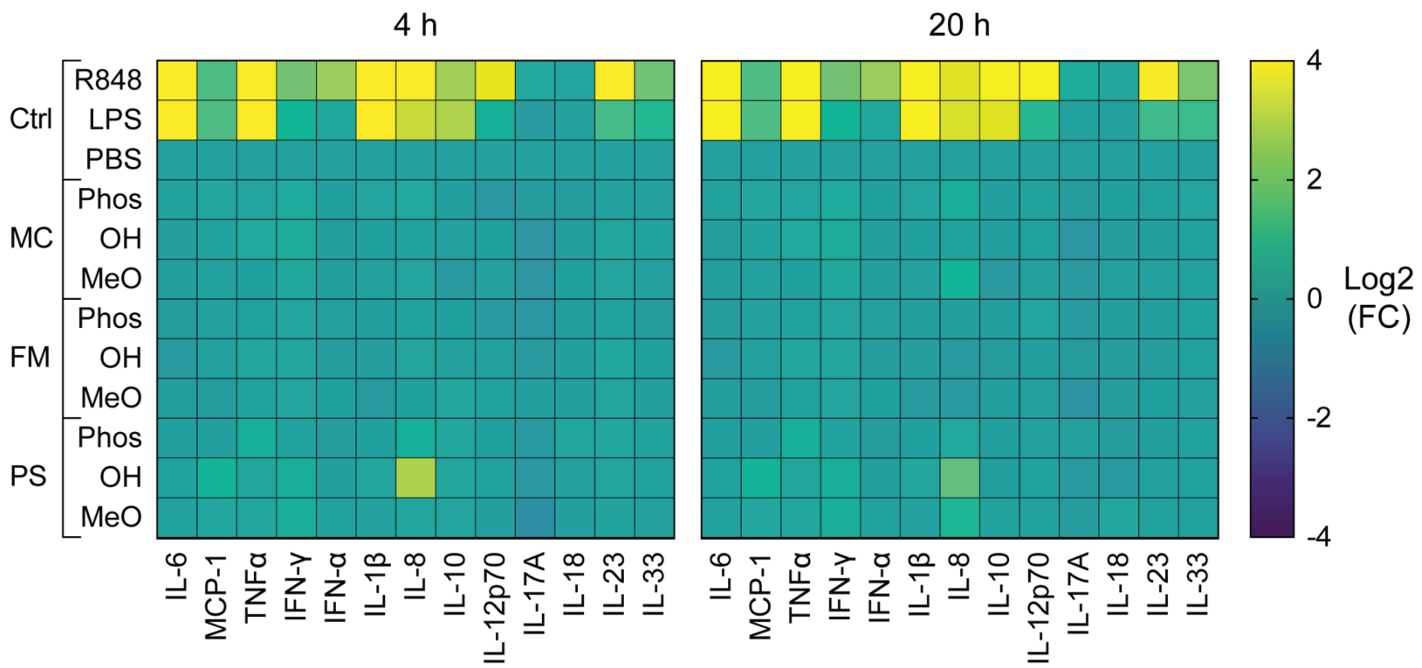


**Figure S17. Average cytokine secretion by PBMCs at 4 h and 20 h.** Averages (n = 3) are displayed from all three donors (individual donor data is displayed in Figure S16).

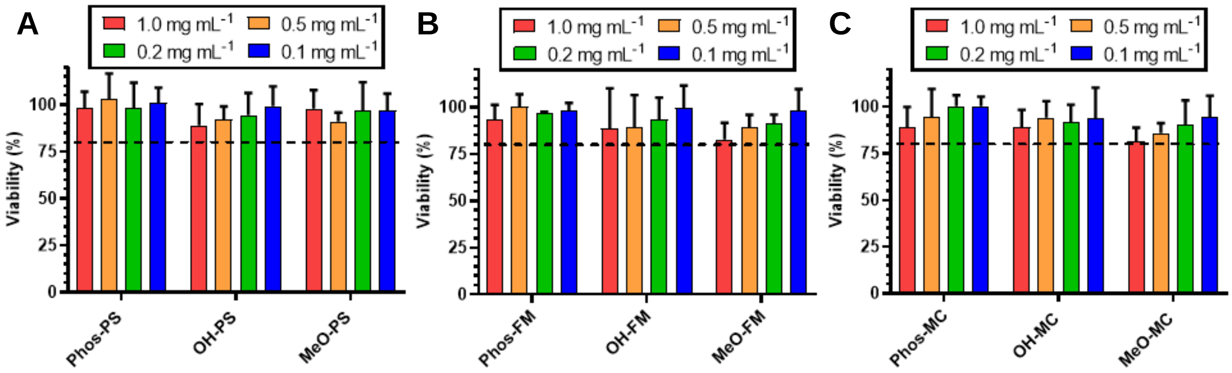


**Figure S18. Viability assessment of nanocarrier formulations with THP-1 monocytes.** Impact on cell viability for Phos-, OH-, and MeO-functionalized (**A**) PSs, (**B**) FMs, and (**C**) MCs (n = 8). Error bars represent s.d. Cell viability was determined through the following formula: % cell viability = (OD of nanocarrier treated sample/OD of untreated sample) × 100.

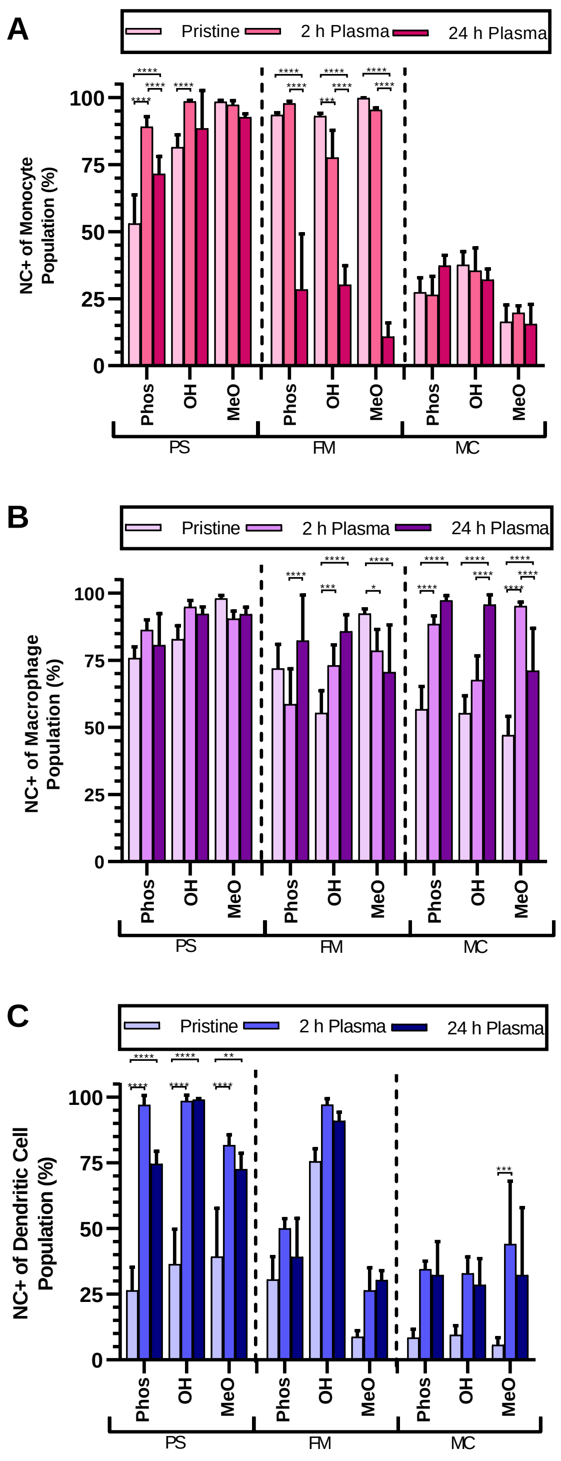


**Figure S19. Statistical significance of differences in cellular uptake of nanocarriers.** The flow cytometric analysis of nanocarrier uptake by (**A**) THP-1 monocytes (n = 10; exception: OH PS with 24 h plasma, n = 9), (**B**) THP-1 differentiated macrophages (n = 10), and (**C**) immature dendritic cells (n = 4). Error bars represent s.d. Statistical significance was determined using Tukey's multiple comparison test, with a 5% significance level. **p*<0.05, ***p*<0.01, ****p*<0.001, *****p*<0.0001. Error bars represent s.d. Cellular uptake data is re-displayed from **Fig. 6B-D**, as bar plots, for the purpose of comparing statistical significance.

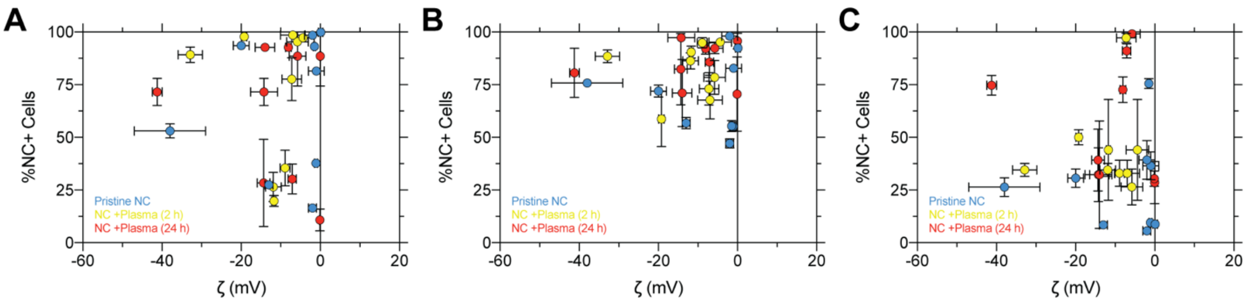


**Figure S20. Relationship between nanocarrier zeta potential and cellular uptake.** The %NC+ cells and nanocarrier zeta potential is shown for uptake data from (**A**) monocytes, (**B**) macrophages, and (**C**) dendritic cells. Pristine (blue), 2 h incubation with human plasma (yellow), and 24 h incubation with human plasma (red) are displayed as groupings on each plot. Spearman r: monocytes (pristine r: 0.29, 2h r: 0.31, 24h r: 0.59); macrophages (pristine r: 0.13, 2h r: 0.18, 24h r: 0.17); dendritic cells (pristine r: 0.58, 2h r: -0.24, 24h r: -0.28). All spearman correlations, calculated within groupings, were not statistically significant (*p*>0.05 in all cases).

### Supplementary Tables

**Table S1. Physicochemical characteristics of spherical and cylindrical nanocarriers.**

| Nanocarrier | | Hydrodynamic Diameter† (nm) | PDI†† | Cross Sectional Diameter‡ (nm) | Contour Length‡ (µm) | Zeta Potential (mV) |
|---|---|---|---|---|---|---|
| Phos | MC | 19 ± 5 | 0.11 ± 0.03 | - | - | -13.0 ± 1.0 |
| OH | MC | 17 ± 3 | 0.08 ± 0.02 | - | - | -1.1 ± 0.4 |
| MeO | MC | 16 ± 5 | 0.07 ± 0.01 | - | - | -2.0 ± 1.0 |
| Phos | FM | - | - | 34 | 2.5 | -20.0 ± 2.0 |
| OH | FM | - | - | 36 | 1.2 | -1.5 ± 0.3 |
| MeO | FM | - | - | 36 | 2.5 | 0.1 ± 0.6 |
| Phos | PS | 55 ± 4 | 0.10 ± 0.01 | - | - | -38.0 ± 9.0 |
| OH | PS | 58 ± 17 | 0.13 ± 0.01 | - | - | -1.0 ± 2.0 |
| MeO | PS | 68 ± 5 | 0.14 ± 0.02 | - | - | -2.0 ± 2.0 |

†Number average value recorded via dynamic light scattering for spherical nanocarriers
††Calculated from number average diameter distribution recorded via dynamic light scattering for spherical nanocarriers
‡Values acquired from model fit of small angle X-ray scattering for cylindrical nanocarriers

**Table S2. Protein-induced changes to nanocarrier zeta potential.**

| Nanocarrier | | †Δζ (2 h) (mV) | †Δζ (24 h) (mV) | ††ΔΔζ (mV) |
|---|---|---|---|---|
| Phos | MC | -0.5 ± 1.2 | 5.7 ± 2.0 | 6.1 ± 2.3 |
| OH | MC | -4.5 ± 1.0 | 6.6 ± 1.2 | 11.1 ± 1.6 |
| MeO | MC | -1.2 ± 1.2 | 9.7 ± 2.7 | 10.8 ± 1.9 |
| Phos | FM | 1.1 ± 0.9 | 9.1 ± 2.1 | 8.1 ± 2.3 |
| OH | FM | -4.4 ± 1.5 | -4.7 ± 0.6 | -0.3 ± 1.6 |
| MeO | FM | 1.7 ± 1.6 | 3.1 ± 1.8 | 1.4 ± 2.4 |
| Phos | PS | 11.4 ± 2.2 | 3.9 ± 1.3 | -7.4 ± 2.5 |
| OH | PS | -3.1 ± 1.9 | 1.4 ± 1.3 | 4.5 ± 2.3 |
| MeO | PS | -7.6 ± 4.2 | -8.1 ± 0.4 | -0.5 ± 4.2 |

†Δζ is calculated as the nanocarrier-protein complex zeta potential minus that of the nanocarrier absent of protein.
††ΔΔζ is calculated as the Δζ24 h minus the Δζ2 h.
In all cases, error represents the s.e.m.

**Table S3. Whole blood patient data.**

| Patient | Gender | Age | Ethnicity |
|---|---|---|---|
| KP #55619 | Female | 33 | African American |
| KP #55613 | Male | 41 | African American |
| KP #55614 | Male | 47 | Caucasian |
